# Amino acid variability, tradeoffs and optimality in human diet

**DOI:** 10.1101/2021.06.16.448627

**Authors:** Ziwei Dai, Jason W. Locasale

**Author notes:** Corresponding author: Jason W. Locasale.

## Abstract

While the quality of fat (e.g. saturated/unsaturated) and carbohydrate (e.g. whole grain/simple sugars) intake has been of great interest, less attention has been made to the type of protein and resulting amino acid intake profiles in human diets. Studies at the molecular level however demonstrate that dietary amino acid intake produces substantial effects on health and disease such as cancer by modulating metabolism. How these effects may manifest in human food consumption and dietary patterns is unknown. We developed a series of algorithms to map, characterize and model the landscape of amino acid content in human food, dietary patterns, and individual consumption including relations to health status, covering over 2,000 foods, ten dietary patterns, and over 30,000 dietary records. We found that the type of amino acids contained in foods and human consumption is highly dynamic with variability far exceeding that of fat and carbohydrate. Some amino acids positively associate with diseases such as obesity while others contained in the same food negatively link to disease. Using linear programming and machine learning, we show that these health trade-offs among can be accounted to satisfy biochemical constraints in food and human eating patterns to construct a Pareto front in dietary practice, a means of achieving optimality in the face of tradeoffs that are commonly considered in economic and evolutionary theories. Thus this study may enable the design of human protein quality intake guidelines based on a quantitative framework.

## Introduction

Diet is generally considered to be a major determinant of human health and disease^1-5^. Numerous dietary recommendations, such as the Dietary Guidelines for Americans^6^, have been developed. These dietary recommendations often focus on two major goals: to increase the diversity and nutrient density of the foods consumed, and to reduce the intake of certain components known to increase risk of disease^7-9^. Such restrictions involve limiting the intake of certain types of carbohydrate and fat such as added sugar, saturated fat and trans-fat, and has rationale based on epidemiology, human^10-12^ and model organism research^13,14^. While it has been widely acknowledged that the types of dietary carbohydrate and fat are important determinants of the quality of a diet, protein the other macronutrient^15^, is often neglected. In most human nutritional studies albeit with exceptions, protein is considered as a single variable and often held constant^16^. Nevertheless, each amino acid has its specific metabolism^17^ and is important for numerous cellular and physiological processes. A growing number of studies shows that variation in dietary intake of amino acids such as serine, glycine, asparagine, histidine, and methionine mediates health and disease including cancer through defined molecular mechanisms^18-28^. Altogether there is a rationale for investigating in a systematic manner amino acid intake in human diets and possible consequences on health.

In this study, we investigated the variability of amino acids in human food and diets and find variability commensurate with what is observed in fats and carbohydrates. Based on optimizing associations with health status, we use these analyses to devise guidelines for dietary amino acids. Finally, we implement machine learning algorithms to design personalized diets based on amino acid intake that correspond to optimality in specified health statuses.

### Amino acid landscape of human food

To characterize the variability of amino acid levels in human food, we first constructed a database consisting of amino acid profiles in three levels of human dietary components: individual foods, dietary patterns or representations of patterns of food consumption (e.g. Western, Mediterranean, Japanese, Keto, etc), and dietary records containing daily reported food intake (Figure 1). The abundance of 18 amino acids in 2,335 foods was collected based on nutritional profiles in the United States of America Department of Agriculture National Nutrient Database for Standard Reference Legacy Release (USDA SR) (Figure 1a, methods). 18 of the 20 amino acids were considered because during quantitation, amino acids which largely exist in protein-bound forms, require hydrolysis into free amino acids during which amino groups from glutamine and asparagine are also hydrolyzed to make glutamic and aspartic acid. Thus, the abundance of glutamic acid and aspartic acid from measurements of free amino acid levels reflects the total abundance of glutamate and glutamine, and the total abundance of aspartate and asparagine, respectively. The distributions of amino acid abundance over 2,335 foods show that each amino acid has considerable variability across foods (Coefficient of variation > 0.2 for all amino acids, Figure 1b), and amino acids most abundant in human food are glutamine/glutamate (median = 0.16 g/g total amino acids), asparagine/aspartate (median = 0.095 g/g total amino acids), leucine (median = 0.082 g/g total amino acids), and lysine (median = 0.076 g/g total amino acids). On the other hand, amino acids with the lowest abundance in human foods are cystine (median = 0.012 g/g total amino acids), tryptophan (median = 0.012 g/g total amino acids), methionine (median = 0.024 g/g total amino acids), and histidine (median = 0.028 g/g total amino acids). This ordering largely resembles the abundance of amino acids in the proteomes which are conserved across living organisms^29,30^. Principal component analysis (PCA) shows that amino acid abundances can be clustered by different categories of foods (Figure 1c, d, methods). Highly variable amino acids include those whose dietary modulation has molecular links to cancer progression and health outcomes, such as methionine (0.031 g/g total amino acids in eggs compared to 0.013 in legumes) and serine (0.076 g/g total amino acids in eggs compared to 0.039 in lamb, veal, and game meat). To quantify the variability of amino acid abundance across foods, we computed the F-statistic from one-way analysis of variance (ANOVA), and compared the resulting F-statistic values with those of carbohydrates (i.e. dietary fiber and sugar) and fats (i.e. saturated fat, monounsaturated fat, and polyunsaturated fat). Notably, we found that the ANOVA F-statistics for amino acids were comparable to or higher than those for carbohydrates and fats (Figure 1e, methods), especially for the amino acids methionine, histidine, lysine, and proline (F-statistic = 816.2 for methionine, 566.1 for histidine, 504.3 for lysine, and 362.9 for proline compared to the range of 45.0 to 119.6 for carbohydrates and the range of 125.2 to 746.3 for fats, Figure 1e, f), highlighting the variability of amino acid abundance in foods which has been largely overlooked previously. Taken together, these results suggest that differences in food intake due to the high variability in amino acid content may lead to differences physiological and cellular effects on metabolism.

**Figure 1.**
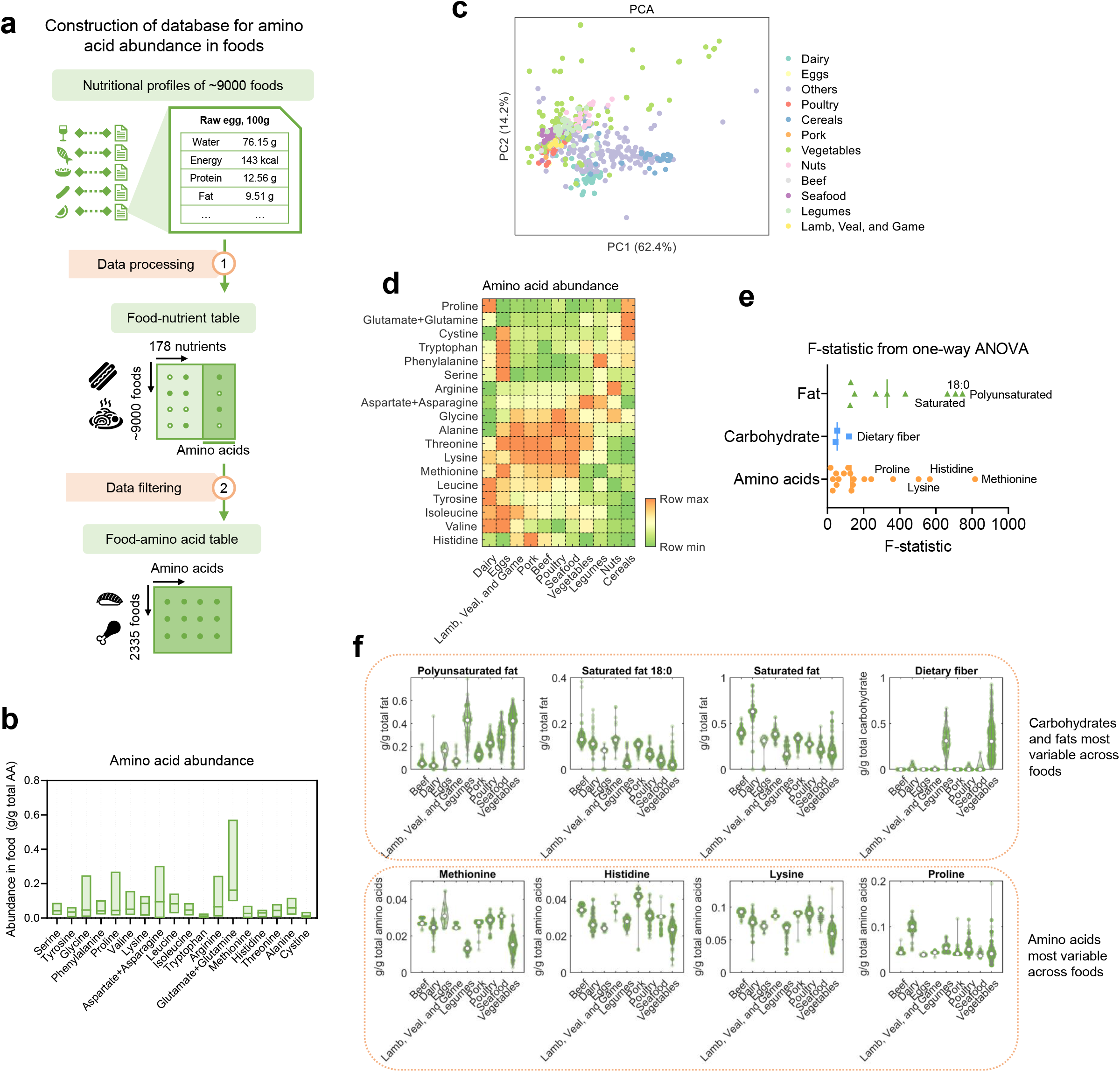
Amino acid landscape of human foods. a. Workflow for construction of the database for amino acid abundances in human foods. b. Ranges of amino acid abundance in human foods. The horizontal lines indicate median values. c. Principal Components Analysis (PCA) of amino acid profiles in human foods. Each dot represents a food. Colors of the dots indicate different categories of the foods. d. Average amino acid abundance in different categories of human foods. e. F-statistic values from one-way ANOVA comparing abundance of amino acids, different types of carbohydrate, and different types of fat across human foods. f. Violin plots showing the distributions of abundance of amino acids, carbohydrates, and fats that are the most variable across human foods. The circles indicate median values. Green dots indicate individual values.

### Human dietary patterns are variable in amino acid content

Dietary patterns can be grouped according to eating patterns that often have a cultural or societal element. They can be characterized by a combination of certain types of foods consumed (e.g. Mediterranean diet, which includes high amounts of plant-based foods, high to moderate amounts of seafood, low consumption of red meat, and olive oil as the main source of added fat^31^), or a specific intake profile of certain nutrients (e.g. ketogenic diet, which is defined by very high intake of fat and very low intake of carbohydrate). Adherence to certain dietary patterns, such as the Mediterranean diet or Japanese diet, has been associated with increased lifespan and lower risk of disease^32-34^. Moreover, some emerging dietary patterns, such as the ketogenic diet and the Paleo diet, have recently been shown in some settings to have benefits on metabolic health, neural function, and longevity^35-38^. However, it is unclear whether these dietary patterns differ in their amino acid content, and whether the variability in amino acid abundance across dietary patterns contributes to the health outcomes associated with these diets.

To further understand the relationship between human dietary patterns and amino acid intake, we next developed an algorithm to quantitatively evaluate amino acid abundance in ten representative human dietary patterns (Figure 2a, S1, Supplementary Methods). Among these dietary patterns, the Mediterranean diet and Japanese diet are two traditional diets believed to have beneficial influences on health, while the Dietary Approaches to Stop Hypertension (DASH) diet consists of consumption of a variety of low-fat and minimally processed foods, and the American diet, which represents the dietary behaviors of a typical individual in western society is also considered. We also include diets that restrict the consumption of certain foods (Paleo diet, vegetarian diet, plant-based diet), diets limiting carbohydrate intake (ketogenic diet, Atkins diet), and a USDA recommended diet defined based on the daily nutrient intake goals in the USDA 2015-2020 dietary guidelines for Americans^6^. We first computed the range of amino acid intake (i.e. grams of each amino acid consumed per day) for each dietary pattern using a linear programming algorithm we developed (Figure 2b, Supplementary Methods) and found that, although none of these dietary patterns includes any constraint on amino acid intake, they still differ greatly with each other in the values of amino acid consumption. Moreover, each dietary pattern allowed for substantial flexibility in the intake of all amino acids (maximal daily intake/minimal daily intake > 20 for all dietary patterns and amino acids, Figure 2b), revealing the possibility to modulate amino acid intake under a certain dietary pattern.

**Figure 2.**
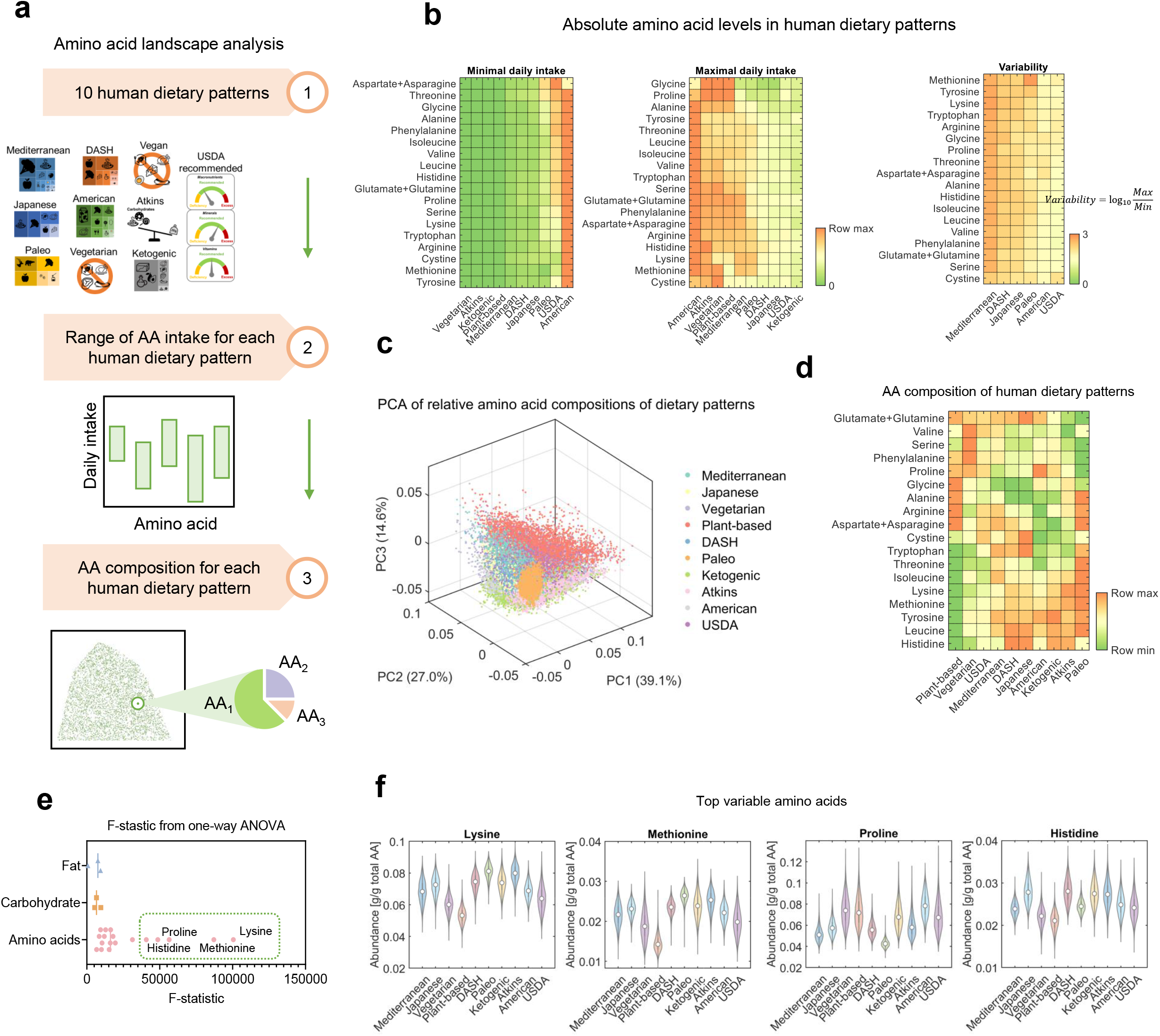
Amino acid landscape of human diets. a. Workflow for the computational modeling of amino acid abundance in human dietary patterns. b. Absolute levels of amino acids in human dietary patterns quantified by the minimal and maximal daily intake values of amino acids in each dietary pattern. c. PCA of relative amino acid compositions of human diets sampled for all ten dietary patterns. Each dot represents for a diet. Colors of the dots indicate different dietary patterns. d. Average amino acid composition of the ten human dietary patterns. e. F-statistic values from one-way ANOVA comparing the composition of amino acids, carbohydrates, and fats across human dietary patterns. f. Violin plots showing the distributions of amino acids that are the most variable across human dietary patterns. The circles indicate median values.

To quantify the variability of amino acid composition that is independent of energy and protein intake, we developed a sampling algorithm based on the accelerated convergence hit-and-run method^39^ to quantify the amino acid composition of each diet by sampling 50,000 instances of each diet (Supplementary Methods). We first confirmed that the sample size of 50,000 was sufficient to capture the distribution of amino acid abundance in a dietary pattern based on the convergence of the sample mean and standard deviation values (Figure S2a). PCA of the sampled diets (Figure 2c) and comparison of mean values (Figure 2d) showed that the ten dietary patterns also have different signatures of amino acid composition. Notably, differences in amino acid composition also exists between dietary patterns similar to each other such as the vegetarian diet and plant-based diet. Indeed, we observed a 30% of difference in methionine abundance between vegetarian diet and plant-based diet (0.019 g methionine/g total AAs in vegetarian diet compared to 0.014 in plant-based diet), suggesting that small changes in the choice of foods result in substantial differences in amino acid intake (Figure 2d, Figure S2b). We also estimated compositions of carbohydrates and fats in these diets (Figure S2b), and quantified the variability of amino acid composition across human diets using F-statistic values from one-way ANOVA, and compared it with the variability of carbohydrates and fats across dietary patterns (Figure 2e). Strikingly, we found that the variability of amino acid composition across diets was much higher than that of carbohydrates and fats, with the amino acids lysine, methionine, proline and histidine being the most highly variable across human dietary patterns (F-statistic > 50,000 compared to less than 10,000 for carbohydrates and fats, Figure 2e-f, S2b-c). Among these amino acids, lysine, histidine and methionine are significantly lower in instances of the plant-based diet, and proline is significantly lower in Paleo diet (Figure 2f). The amino acid signatures of human dietary patterns were further validated by measurements of fasting blood concentrations of the amino acids leucine, isoleucine, and alanine in human subjects eating plant-based or ketogenic diet (Figure S2d)^40^. Taken together, these results reveal that the biggest difference in macronutrient composition across human dietary patterns is in amino acid content, and not that of carbohydrates or fats. How the diversity in dietary amino acids results in different health outcomes remains an open question, which may begin to be answered with nutritional and health data in large populations of humans.

### Landscape of amino acid intake in human dietary records

Next, we considered individual dietary amino acid intake records across a population of individuals from diverse ethnic and cultural backgrounds. We reconstructed the dietary amino acid intake profiles in more than 30,000 human subjects in the United States based on dietary records in the National Health and Nutrition Examination Survey (NHANES) 2007-2014 datasets (Figure 3a). Since the NHANES datasets do not direct include dietary amino acid intake values, we developed a set of computational tools for data imputation and mapping to reconstruct the amino acid profiles for the dietary records based on two additional datasets, the USDA SR food nutritional database and the Food and Nutrient Database for Dietary Studies (FNDDS) (Figure 3a). Data imputation using random forest (RF) regression, which outperformed other methods in the accuracy of imputation (Figure S3a, b), was applied to estimate the missing values of amino acid levels in the USDA SR dataset. The imputed datasets were then used to construct amino acid profiles for the FNDDS and NHANES records by mapping foods in the USDA dataset to foods in the FNDDS dataset which were then used to compute nutrient intake values in the NHANES dietary records (Figure 3a, Supplementary Methods). To assess the limitations of self-reported dietary records in the NHANES data, we compared our computed nutrient intake values with measurements of blood concentrations of related metabolites such as Vitamin D (Figure 3b). Next, to validate the reconstructed amino acid intake levels, we first compared the total intake of amino acids and intake of protein in each dietary record and confirmed that the reconstructed total amino acid intake closely resembles the known total protein intake (Pearson correlation = 0.99, p-value < 10^−323^, Figure 3c), concentrations of amino acids in human blood (Spearman correlation = 0.52, p-value = 0.03, Figure 3d), uptake fluxes of amino acids in human cell lines, which reflect demands of amino acids in cultured human cells (Spearman correlation = 0.70, p-value = 0.01, Figure 3e), and amino acid composition of several culture mediums (Spearman correlation > 0.5 and p-value < 0.05 for 4 out of 7 culture media, Figure S3c). The high correlation between dietary amino acid intake and physiological parameters related to amino acids suggests that our reconstructed amino acid intake data may reflect some aspects of physiological metabolism, and suggest that the cellular behaviors and tissue microenvironment in amino acid metabolism reflect to some extent dietary intake of amino acids despite the many other factors that influence cellular metabolism.

**Figure 3.**
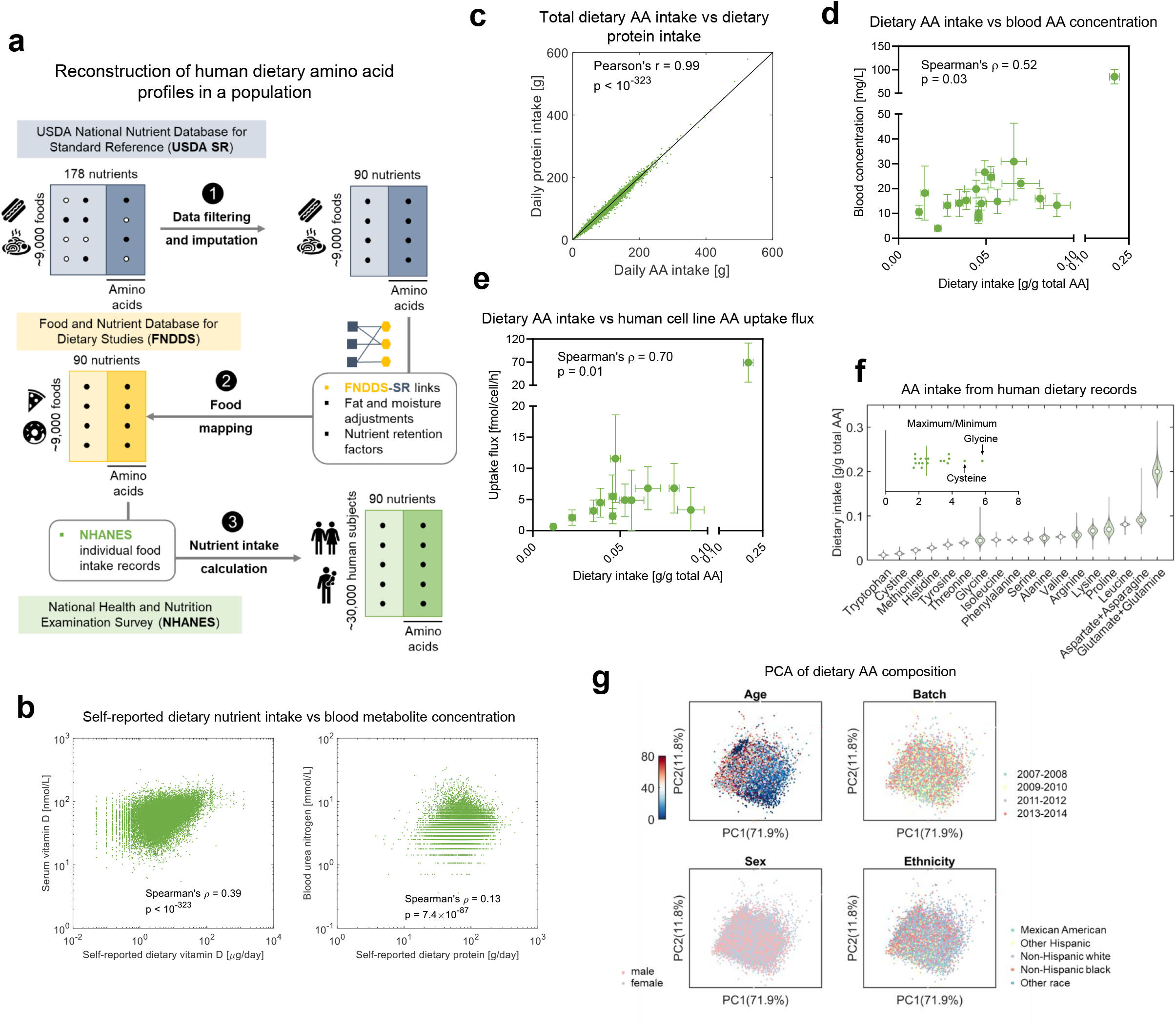
Landscape of human dietary amino acid intake. a. Workflow for reconstruction of the database consisting of amino acid intake profiles in human dietary records. b. Comparison between nutrient intake values in the self-reported dietary records and laboratory measurements of nutrient-related metabolites in blood. c. Comparison between total dietary amino acid intake in the reconstructed amino acid intake database and dietary protein intake in the original dietary records. d. Comparison of the reconstructed human dietary amino acid intake values to blood concentrations of amino acids. The dots represent for mean values and error bars for standard deviations. e. Comparison of the reconstructed human dietary amino acid intake values to uptake fluxes of amino acids. The dots represent for mean values and error bars for standard deviations. f. Distributions of amino acid intake in human dietary records. The circles indicate median values. g. PCA analysis of amino acid intake values in human dietary records showing their association with age, sex, ethnicity, and batch of the data.

We then evaluated the overall variability in the intake of each amino acid based on the ratio of maximal to minimal intake values in the human dietary records (Figure 3f), and performed PCA on the reconstructed dietary amino acid profiles to report the association between dietary amino acid composition and demographic variables such as age, sex, and ethnicity (Figure 3g). We found that among the population included in the NHANES 2007-2014 cohorts, daily intake of amino acids typically varies by two to six fold (e.g. maximal intake/minimal intake = 4 for tryptophan, 2.5 for methionine, 6.2 for glycine, and so on). Dietary amino acid composition profiles showed no difference between batches (Figure 3e), thus confirming that our reconstruction is not biased by batch effect. Interestingly, dietary intake of amino acids was found to correlate with age, while no dependency on other demographic variables such as sex and ethnicity was observed (Fig 3e, Figure S4). These reconstructed dietary amino acid intake profiles allow us to examine the quantitative relationship between dietary amino acids and human health.

### Dietary amino acid intake associations with human health

We next attempted to link dietary amino acid intake and incidence of several human diseases based on the reconstructed dietary amino acid intake profiles and clinical records available in the NHANES database. We focused on chronic diseases that are a major concern to human health such as cardiovascular disease, diabetes, and cancer. We retrieved the medical records of 18,196 adult subjects in the NHANES 2007-2014 datasets and defined quantitative scores describing the incidences of hypertension, obesity, cancer, and diabetes based on the examination, laboratory, and questionnaire datasets (Figure 4a, Methods). We first computed partial Spearman’s rank correlation coefficients as a metric to evaluate the association between dietary amino acid composition and the incidences of the four diseases while controlling for confounders including demographic and lifestyle-related factors (Supplementary Figure 5). We identified many amino acid intake-disease associations involving all four diseases considered (statistically significant associations in 21 out of 72 amino acid-disease pairs, Figure 4b, methods), among which obesity showed the strongest association with dietary amino acid composition (obesity incidence positively correlated with the intake of threonine, histidine, alanine, glycine, lysine and methionine, and negatively correlated with intake of tryptophan, phenylalanine, valine, serine, asparagine, aspartate, glutamine, and glutamate, Figure 4b). These associations between dietary amino acid intake and obesity were consistent with some observations in molecular studies, such as the anti-obesity functions of dietary tryptophan and pro-obesity functions of methionine in mice^41,42^. As a control, we also correlated the incidence of the four diseases with dietary intake of different types of carbohydrates and fats. Counterintuitively, we found much fewer statistically significant associations between dietary intake of carbohydrate and fat (9 significant associations out of 40 disease-nutrient pairs, Figure 4c). These results together highlight the unexpectedly strong association between that dietary intake of amino acids and human disease which exceeds the association for dietary carbohydrates and fats. To further explore these questions, we performed a comparison of the association between nutrients and human health using machine learning models predicting health outcomes from different types of nutritional variables (Figure 4d). We categorized nutritional variables included in the NHANES database into six groups, including energy, macronutrients, macronutrient compositions (i.e. fractions of different types of carbohydrate and fat in total carbohydrate and fat intake), vitamins, minerals, amino acid compositions (i.e. intake of each amino acid with the unit g/g total AA), and other nutrients. For each disease, nutritional variables in each group were used as covariates to build a logistic regression model to predict the incidence of that disease. The area under receiver operating characteristic curve (AUC) with 5-fold cross validation was used to assess the performance of each group of nutritional variables in predicting disease incidence, which reflects strength of the association between dietary intake of those nutrients and that disease. We found that dietary amino acid composition was predictive of incidence of all diseases except for cancer (AUC > 0.5, 5-fold cross validation), and achieved accuracy of prediction comparable to or higher than that of dietary carbohydrate and fat intake for obesity and hypertension (AUC = 0.55 for amino acids compared to 0.55 for macronutrient composition in predicting obesity, and AUC = 0.53 for amino acids compared to 0.52 for macronutrient composition in predicting hypertension, Figure 4e). The reason that dietary amino acid intake was unable to predict cancer outcome was probably for the reason that different types of cancers were not distinguished in the analysis, the population included remissions, and the frequency of cancer in the dataset is relatively low (1844 cases out of 18469 individuals). Nevertheless, the higher accuracy of amino acid intake in predicting obesity and hypertension incidence in humans provides a rationale for optimization of dietary amino acid intake.

**Figure 4.**
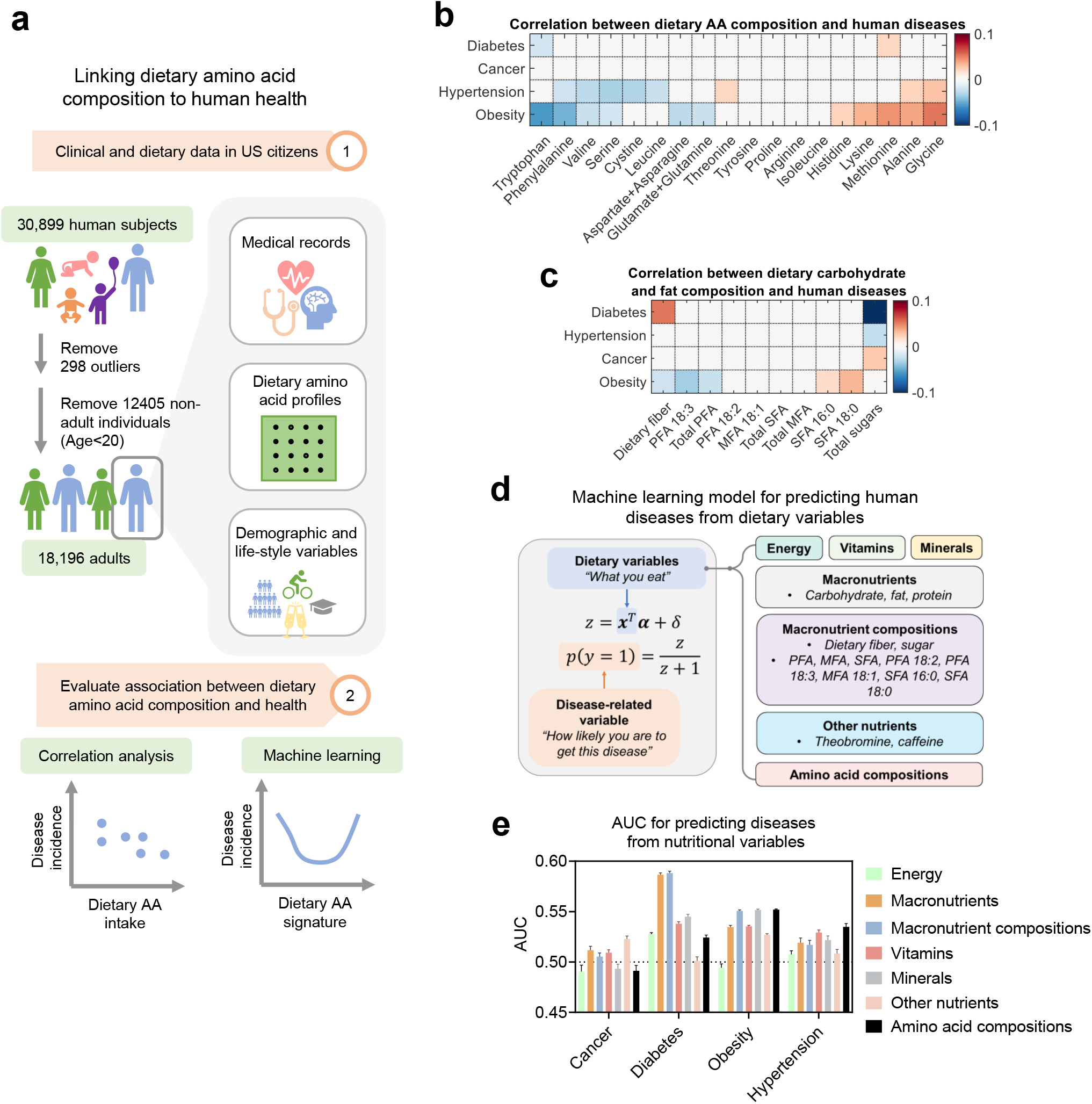
Amino acid intake is predictive of human health. a. Workflow for the analysis of association between dietary amino acid intake and human health. b. Partial Spearman correlation between incidences of human diseases and dietary intake of amino acids. c. Partial Spearman correlation between incidences of human diseases and dietary intake of different types of carbohydrate and fat. d. Framework of the machine learning model predicting incidence of human diseases from different groups of dietary variables. e. AUC values for predicting incidence of human diseases from different groups of dietary variables. Error bars indicate standard deviations.

### Guidelines for dietary amino acids and diet design

Dietary recommendations, such as these in the USDA Dietary Guidelines for Americans, often involve suggestions to consume a variety of minimally processed foods and recommended ranges for intake of nutrients including macronutrients, vitamins, and minerals. Since dietary intake of amino acids has been associated with health outcomes both in molecular studies and by our analysis thus far, we sought to develop an Artificial Intelligence (AI) -based approach for identification of dietary guidelines for amino acids and design of personalized human diets optimizing their amino acid composition.

First, we developed an algorithm for identification of amino acid intake guidelines based on the associations between dietary amino acid intake and human health (Figure 5a). We focused on obesity since it had the highest incidence and was found to have the strongest association with dietary amino acid intake among the four diseases considered in this study (Figure 4b). We classified obesity-associated amino acids into three categories (Figure 5b), including amino acids for which the intake positively associate with obesity incidence (‘positive association’), negatively associate with obesity incidence (‘negative association’), or associate with obesity incidence with a non-monotonic, U-shaped relationship (‘U-shaped relationship’). The amino acids phenylalanine, aspartate/asparagine, tryptophan, valine and glutamate/glutamine fell into the negative association group. On the other hand, the amino acids glycine, alanine, methionine, lysine, histidine were categorized into the positive association group. The association between intake of dietary amino acids and obesity was not due to changes in calorie intake, since amino acids positively associated with obesity were either negatively or positively correlated to calorie intake, and *vice versa* (Figure S6a).

**Figure 5.**
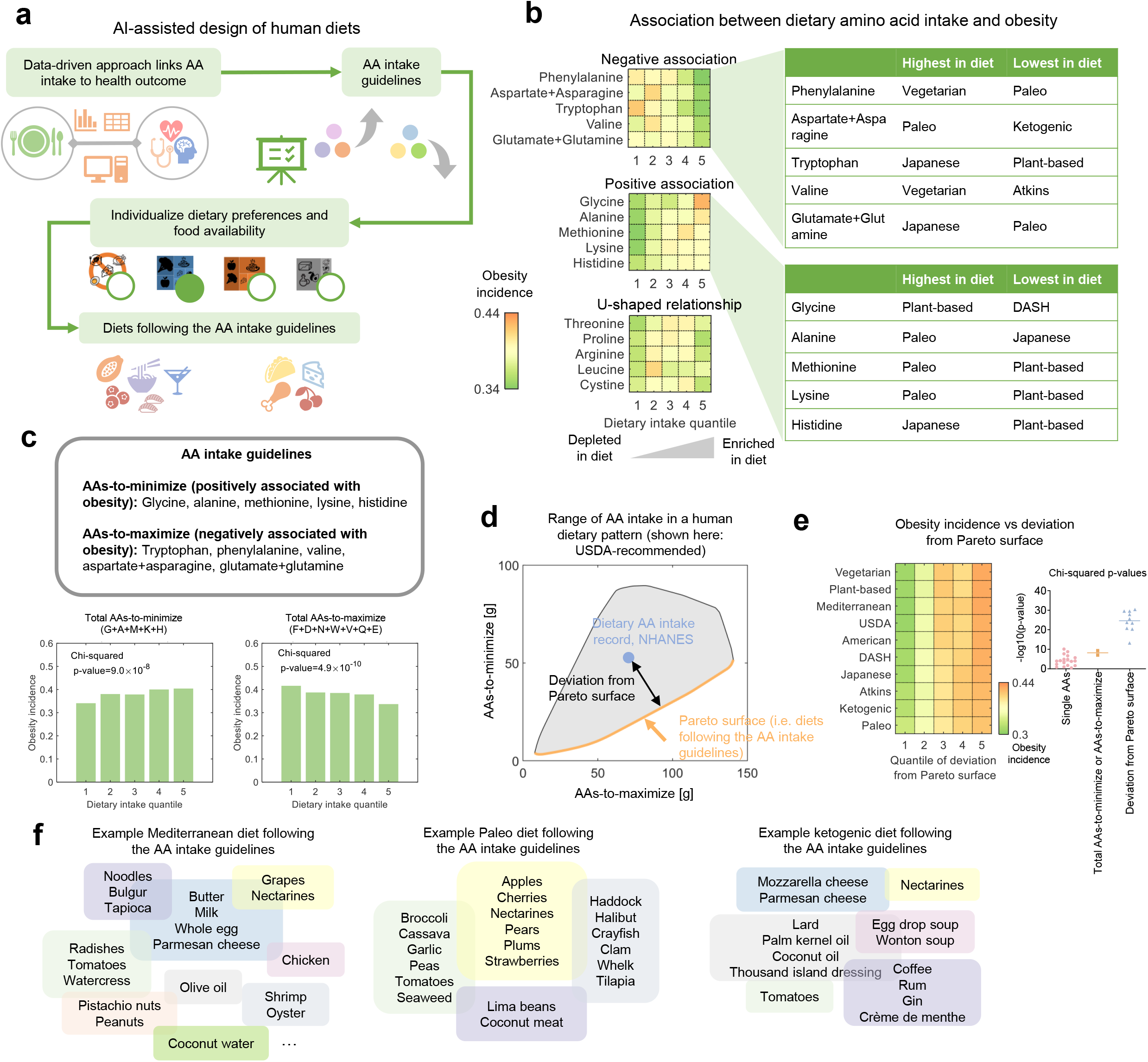
AI for dietary amino acid guidelines and personalized diet design. a. Workflow for AI-assisted identification of dietary amino acid guidelines and design of personalized diets. b. Three types of association between dietary amino acid intake and obesity in humans. c. Identification and confirmation of amino acid intake guidelines based on the association between dietary amino acids and obesity. d. Ranges of intake of total amino-acids-to-maximize and amino-acids-to-minimize in the dietary pattern of USDA-recommended diet (grey shaded region) and the Pareto surface (orange bold curve) corresponding to the two guidelines, i.e. maximizing total amino-acids-to-maximize, and minimizing total amino-acids-to-minimize. e. Associations between the obesity incidence and deviation of dietary records from the Pareto surface. Chi-squared p-values were computed to assess the significance levels of the associations. f. Examples of diets designed according to the amino acid intake guidelines and personalized preferences of dietary patterns.

We also examined whether there exists a dietary pattern that can minimize the intake of the amino acids positively associated with obesity while maximizing the intake of the amino acids negatively associated with obesity. To our surprise, no dietary pattern was able to satisfy all of these requirements. For instance, the Paleo diet has the highest levels of aspartate and asparagine, which negatively associate with obesity. Nevertheless, the Paleo diet also has the highest intake of alanine, methionine and lysine, which all positively associate with obesity incidence. These results reveal the complexity in the relationship between dietary amino acid intake and obesity, indicating trade-offs between the goals of maximizing or minimizing different groups of amino acids which should be considered while designing dietary guidelines for amino acids.

We therefore sought to define dietary amino acid intake guidelines based on the association between dietary amino acids and obesity (Figure 5c), that is, to minimize the total intake of amino acids that positively associate with obesity (i.e. AAs-to-minimize, including glycine, alanine, methionine, lysine, histidine), and to maximize the total intake of amino acids that negatively associate with obesity (i.e. AAs-to-maximize, including tryptophan, phenylalanine, valine, aspartate+asparagine, glutamate+glutamine). We first confirmed that both total AAs-to-minimize and total AAs-to-maximize were significantly associated with obesity incidence (Chi-squared p-value = 9.0×10^−8^ for total AAs-to-minimize and 4.9×10^−10^ for total AAs-to-maximize, Figure 5c).

We then further characterized the trade-off between the requirements of minimizing total AAs-to-minimize and maximizing total AAs-to-maximize by constructing the Pareto surface based on the two requirements (Figure 5d). The concept of Pareto optimality has been widely applied in economics and engineering, and introduced to biology to characterize the trade-off between multiple tasks of bacteria, cancer cells, and organisms^43-46^. For each dietary pattern, there exists a Pareto surface consisting of diets that best balance the needs to minimize total AAs-to-minimize and to maximize total AAs-to-maximize, meaning that for a diet within the Pareto surface, any other diet following this dietary pattern would never have both higher total intake of AAs-to-maximize and lower total intake of AAs-to-minimize at the same time. We hence developed an algorithm to construct the Pareto surface for each of the ten dietary patterns considered in this study (Figure 5d, S6b, Methods), and quantified the extent by which a specific diet satisfies the two requirements of maximizing total AAs-to-maximize and minimizing total AAs-to-minimize using the deviation from Pareto surface (Figure 5d). For each dietary pattern, we computed the deviation of each NHANES dietary record from its Pareto surface, and found that the deviation from the Pareto surface strongly correlates with obesity incidence (Chi-squared p-values < 10^−10^ for all dietary patterns), implying that diets on the Pareto surface of each dietary pattern are associated with lower risk of obesity. On average, an individual that eats a diet that is the top 20% furthest away from the Pareto surface has a 34% higher chance of being obese compared to one eating a diet among the 20% closest to the Pareto surface (Figure 5e).

These findings not only reveal novel relationship between dietary amino acid intake and health, but also allow us to design diets that have amino acid profiles associated with lower risk of obesity and satisfy personalized needs and requirements such as preferred dietary patterns according to the constructed Pareto surface of the preferred dietary pattern. Hence, based on such strategy, we developed an AI for designing diets including the Mediterranean, Paleo, and ketogenic diet (Figure 5f). Each diet contains a variety of foods from diverse sources and keeps the features of the corresponding dietary pattern.

## Discussion

This study develops data resources and computational techniques to begin to address two major limitations in the nutritional sciences: 1) the lack of systematic collections of nutritional information and 2) the lack of computational tools to probe the connections in food, dietary patterns and practices, and health status. Consequentially, we made a number of findings about the variability of amino acids across different types of human foods and dietary patterns and the unexpected associations between dietary amino acid intake, food and dietary patterns, and health. Unexpected links from amino acid intake to pathology such as obesity highlight non-intuitive diet-disease associations and inherent tradeoffs in amino acid content in food.

While we were able to use the tools we devised to study and make discoveries about the landscape of amino acid intake, these capabilities are generalizable to any systematic analysis of human food and diet. For instance, it is still unclear how dietary patterns and human dietary records differ with each other in micronutrients such as vitamins, minerals, dietary fiber, added sugars, and how personalized diets can be designed to cover more nutritional goals. Application of the algorithms we developed in this study may help address these questions.

This study has some limitations. First, the association between dietary amino acids and human diseases is observational and does not directly imply causality. Nevertheless, some amino acid-disease associations identified by our analysis have been observed in experimental studies. For instance, tryptophan, which was found to be negatively associated with obesity in our study, was shown in mice to reduce appetite and weight gain through the production of serotonin in brain^41^. On the other hand, dietary restriction of methionine in mice and human has been shown to improve metabolic health and increase fat oxidation, which may contribute to the anti-obesity effects of dietary methionine restriction^42,47,48^. Further studies, such as randomized controlled trials that directly compare the health outcomes of diets differing with each other in amino acids, are necessary but also limited to the cohort in consideration and the pre-determined end points.

We also note that the datasets used in this study are not completely free of bias. The majority of records in the databases of foods and human dietary records are western, while foods frequently consumed in other geographical regions and by other cultural groups, such as Asians and Africans, are largely underrepresented. Therefore, application of our findings to non-western populations may be limited. Nevertheless, we are optimistic that this limitation could be addressed by extending the coverage of the existing nutritional and epidemiological datasets to non-western populations^49,50^.

## Methods

### Computer algorithms and their implementation

Details about the computer algorithms used in this study, including these for reconstruction of amino acid landscape in human foods, dietary patterns, and dietary records, are explained in the Supplementary Methods. The algorithms for imputation and reconstruction of amino acid profiles in the NHANES database, including imputation of missing data, and mapping of foods in the USDA SR, FNDDS, and NHANES databases, were implemented in R. All other algorithms used in this study were implemented in MATLAB. The database for amino acid abundance in human foods, dietary patterns and dietary records was implemented in both Microsoft Access database file and Microsoft Excel files. All database files are freely available for download at the GitHub repository: https://github.com/ziweidai/AA_human_diet/tree/main/6-Database.

### Statistical analysis

Principal component analysis was performed using the MATLAB built-in function ‘pca()’. One-way ANOVA was performed using the MATLAB built-in function ‘anova1()’. Logistic regression models were constructed, trained, and evaluated using the MATLAB built-in functions ‘glmfit()’, ‘glmval()’, and ‘perfcurve()’. Chi-squared test was performed using the MATLAB built-in function ‘crosstab()’. Relationships with p-value < 0.05 were considered significant. Partial Spearman’s rank correlation coefficients were computed using the MATLAB built-in function ‘partialcorr()’ with p-values adjusted using the Benjamini-Hochberg procedure. Associations with adjusted p-value < 0.05 were considered significant. Average amino acid abundances in food categories or dietary patterns were computed using the mean values of amino acid abundances across all foods in that food category or instances in that dietary pattern.

## Data and code availability

All code, scripts, and datasets used or generated in this study are available at the GitHub page of Ziwei Dai: https://github.com/ziweidai/AA_human_diet.

## Acknowledgments

The authors thank all members of the Locasale Lab, especially Dr. Zhengtao Xiao, Shiyu Liu, and Yudong Sun, for helpful discussions. JWL thanks the Marc Lustgarten Foundation, the National Institutes of Health (R01CA193256 to JWL), and the American Cancer Society 129832-RSG-16-214-01-TBE for their generous support. Support for computational resources from the Duke Compute Cluster and Data Commons Storage is gratefully acknowledged.

## Author contributions

Z.D. and J.W.L. designed the study, wrote and edited the paper. Z.D. developed the algorithms and analyzed the data.

