## Supplementary Methods and Figures for "Amino acid variability, tradeoffs and optimality in human diet"

### 1. Mathematical models of diets

#### 1.1 Mathematical definition of diets

A diet is defined as a combination of foods consumed by an individual on a daily basis, and the consumed amount of each food included in this diet. A subset of foods (2335 foods in total) in the USDA standard reference food composition database release 28 (USDA SR28) was considered in defining diets. Other foods were discarded in this analysis due to missing values in important nutrients (i.e. carbohydrate, fat, protein, vitamins, minerals, amino acids, and other nutrients whose daily intake values are considered in the USDA 2015-2020 dietary guidelines). Thus, each diet is defined as a numeric vector with 2335 elements, each of which describes the amount of the corresponding food in this diet:

$$\boldsymbol{x} = \begin{bmatrix} x_1 \\ x_2 \\ \vdots \\ x_{2335} \end{bmatrix}$$

For instance, the variable $x_1$ describes the amount (in grams per day) of the first food, which has the identifier '01001, Butter, salted' in the USDA food database, in the diet $\boldsymbol{x}$. If $x_1 = 10$, it means that an individual eating diet $\boldsymbol{x}$ consumes 10 grams of salted butter per day. Since one cannot consume negative amount of foods, all elements of $\boldsymbol{x}$ are nonnegative, i.e. $x_i \geq$

$0, i = 1,2,\cdots,2335$.

#### 1.2 Nutritional values of foods and diets

The nutritional profile of a food is defined by the amount of each nutrient in one gram of the given food. Thus, for each nutrient, there are 2335 values that describe the levels of this nutrient in the 2335 foods. Hence, for each nutrient, we define a 2335-dimensional nutrient abundance vector $\boldsymbol{b}$ that quantifies the abundances of this nutrient in the 2335 foods:

$$\boldsymbol{b} = \begin{bmatrix} b_1 \\ b_2 \\ \vdots \\ b_{2335} \end{bmatrix}$$

The element $b_i$ describes the amount of the corresponding nutrient in one gram of the *i*-th food. For instance, considering the nutrient 'Total lipid (fat)', $b_1$ has the value 0.8111, meaning that each gram of the first food, which has the identifier '01001, Butter, salted', contains 0.8111 gram of fat in total.

Based on the definitions of the diet vector $\boldsymbol{x}$ and nutrient abundance vector $\boldsymbol{b}$ for a nutrient, the total amount of this nutrient in the diet $\boldsymbol{x}$ is $r = \boldsymbol{b}^T\boldsymbol{x}$. Hence, the total amount of a nutrient in a diet is a linear function of amounts of foods in this diet with linear coefficients determined by abundances of this nutrient in different foods.

We next consider the abundances of *m* different nutrients at the same time. For the *i*-th nutrient, let $\boldsymbol{b}_i$ denote the nutrient abundance vector for this nutrient, we can then compute the amounts of nutrients $1,2,\cdots,m$ in the diet $\boldsymbol{x}$:

$$r_i = \boldsymbol{b}_i^T\boldsymbol{x}$$

Thus, the nutritional composition $\boldsymbol{r}$ of the diet $\boldsymbol{x}$ can be written in the following form:

$$\boldsymbol{r} = \begin{bmatrix} r_1 \\ r_2 \\ \vdots \\ r_m \end{bmatrix} = \begin{bmatrix} \boldsymbol{b}_1^T\boldsymbol{x} \\ \boldsymbol{b}_2^T\boldsymbol{x} \\ \vdots \\ \boldsymbol{b}_m^T\boldsymbol{x} \end{bmatrix} = [\boldsymbol{b}_1 \quad \boldsymbol{b}_2 \quad \cdots \quad \boldsymbol{b}_m]^T\,\boldsymbol{x}$$

Let $\mathbf{B} = [\boldsymbol{b}_1 \quad \boldsymbol{b}_2 \quad \cdots \quad \boldsymbol{b}_m]^T$, then we have $\boldsymbol{r} = \mathbf{B}\boldsymbol{x}$. The element $B_{ij}$ in the matrix $\mathbf{B}$ denotes the abundance of the *i*-th nutrient in the *j*-th food.

Specifically, we quantify the abundance of amino acids in foods and diets using similar notation. Let $\mathbf{A} = [\boldsymbol{a}_1 \quad \boldsymbol{a}_2 \quad \cdots \quad \boldsymbol{a}_{18}]^T$ denote the abundances of amino acids in the 2335 foods (i.e. columns of $\mathbf{A}$ are nutrient abundance vectors for amino acids), we can compute the amino acid composition $\boldsymbol{s}$ of each diet $\boldsymbol{x}$ by $\boldsymbol{s} = \mathbf{A}\boldsymbol{x}$. The reason that 18 instead of 20 amino acid variables are considered is that the current protocol for quantification of amino acids in foods requires hydrolysis of protein, which breaks protein-bound amino acids into free amino acids, before quantification of the amino acids. During the protein hydrolysis, the amides glutamine and asparagine are converted to glutamic acid and aspartic acid, respectively.

#### 1.3 Mathematical definition of human dietary patterns

Human dietary patterns are representative modes of diets consumed in certain geographical regions, consumed by certain ethnic groups, or recommended by certain dietary guidelines for the goal of improving health. Examples of human dietary patterns include Mediterranean diet, Japanese diet, Paleo diet and plant-based diet, which are defined by consumption of certain combinations of foods, or ketogenic diet and Atkins diet, which are defined by limited intake of certain nutrients. Thus, each human dietary pattern can be defined mathematically as a set of constraints on the composition of foods or nutrients in a diet that falls into this category of dietary pattern.

##### 1.3.1 Constraints on foods

This type of constraint limits the total amount of certain food allowed in a diet. These constraints set either an upper bound or a lower bound of daily consumption of the corresponding food or group of foods.

For example, in a plant-based diet, consumption of animal products, such as meat, seafood, egg and dairy products, is strictly prohibited. Thus, for each food, we can define a binary label indicating whether this food is animal-based or not:

$$d_i = \begin{cases} 1, & \text{if the i-th food is animal based} \\ 0, & \text{otherwise} \end{cases}$$

A diet is a plant-based diet if and only if the total amount of animal-based foods in this diet equals zero, i.e. $\boldsymbol{d}^T\boldsymbol{x} = 0$. In other words, every diet that satisfies this linear constraint is a plant-based diet, hence the linear constraint $\boldsymbol{d}^T\boldsymbol{x} = 0$ gives a definition of plant-based diet.

Another example is the Mediterranean diet, in which 3 to 6 servings of cereals are consumed each day. Similar to the case of plant-based diet, we can define a vector $\boldsymbol{d}$:

$$\boldsymbol{d} = \begin{bmatrix} d_1 \\ d_2 \\ \vdots \\ d_{2335} \end{bmatrix}$$

$$d_i = \begin{cases} \dfrac{1}{w_i}, & \text{if the i-th food is a cereal-based food} \\ 0, & \text{otherwise} \end{cases}$$

In which $w_i$ is the weight of one serving of the $i$-th food. Thus, in a Mediterranean diet, the daily intake of 3 to 6 servings of cereals can be represented by the linear constraint:

$$3 \leq \boldsymbol{d}^T\boldsymbol{x} \leq 6$$

Similar constraints on daily consumption of other groups of foods, such as seafood, fruits, legumes and so on, can also be defined in this way. These constraints together offer a definition of Mediterranean diet (see Fig S2 for a complete list of food groups included in Mediterranean diet). A diet is a Mediterranean diet if and only if it satisfies all of these constraints on allowed numbers of servings of foods.

A general form of this type of constraint can be written as below:

$$\boldsymbol{l}_f \leq \mathbf{D}\boldsymbol{x} \leq \boldsymbol{u}_f$$

In which $\boldsymbol{l}_f$ and $\boldsymbol{u}_f$ are the lower and upper bounds of the total amount of foods in each category whose total daily consumption is constrained in a specific range in this diet.

**1.3.2 Constraints on absolute levels of nutrient intake**

This type of constraint limits the total daily intake of a nutrient in a diet. Similar to the constraints on the amount of foods, these constraints also determine upper or lower bounds of nutrient intake. For instance, in the Atkins diet, which is a carbohydrate-restricted diet, the recommended total daily intake of carbohydrate is no more than 20 grams. As we have discussed in section **1.2**, the total daily intake of carbohydrate in a diet $\boldsymbol{x}$ is $\boldsymbol{c}^T\boldsymbol{x}$, in which $\boldsymbol{c}$ is the vector of carbohydrate contents of foods. Thus, the Atkins diet can be defined by the linear constraint:

$$\boldsymbol{c}^T\boldsymbol{x} \leq 20$$

This type of constraints can be written in the general form below:

$$\boldsymbol{l}_n \leq \mathbf{C}\boldsymbol{x} \leq \boldsymbol{u}_n$$

In which $\boldsymbol{l}_n$ and $\boldsymbol{u}_n$ are the lower and upper bounds of the total daily intake of nutrients related to this diet, and the *i*-th row in the matrix $\mathbf{C}$ is the nutrient abundance vector of the *i*-th of these nutrients.

#### 1.3.3 Constraints on ratios of nutrient intake

This type of constraint determines allowed ranges of ratios between two nutrients. Most of these constraints are about the percentage contribution of a macronutrient, such as carbohydrate, to the total daily intake of calories. For instance, in a ketogenic diet, at least 70% of the total daily calories come from fat, while a very small fraction (e.g. less than 5%) comes from carbohydrate. Let $\boldsymbol{c}$, $\boldsymbol{f}$ and $\boldsymbol{e}$ denote the nutrient abundance vectors for carbohydrate, fat and calories, these constraints on the relationship between intake of fat, carbohydrate and calories in a ketogenic diet can be written as below:

$$\begin{cases} \boldsymbol{c}^T\boldsymbol{x} \leq 0.05\boldsymbol{e}^T\boldsymbol{x} \\ \boldsymbol{f}^T\boldsymbol{x} \geq 0.7\boldsymbol{e}^T\boldsymbol{x} \end{cases}$$

Rearranging the terms at the two ends of the inequalities, we have:

$$[\boldsymbol{c} - 0.05\boldsymbol{e} \quad 0.7\boldsymbol{e} - \boldsymbol{f}]^T\boldsymbol{x} \leq \boldsymbol{0}$$

Thus, a general form of this type of constraints is:

$$\mathbf{E}\boldsymbol{x} \leq \boldsymbol{0}$$

#### 1.3.4 The general mathematical form for definition of a diet

Summarizing the types of constraints in sections **1.3.1**, **1.3.2** and **1.3.3**, we have the general form of the definition of a diet:

$$\begin{cases} \boldsymbol{x} \geq \mathbf{0} \\ \boldsymbol{l}_n \leq \mathbf{C}\boldsymbol{x} \leq \boldsymbol{u}_n \\ \boldsymbol{l}_f \leq \mathbf{D}\boldsymbol{x} \leq \boldsymbol{u}_f \\ \mathbf{E}\boldsymbol{x} \leq \mathbf{0} \end{cases}$$

It is worth noting that all of these constraints are linear.

### 2. Quantification of amino acid levels in diets

The goal of this part of analysis is to quantify abundance of amino acids in human diets and identify quantitative amino acid signatures for each dietary pattern such as Mediterranean, American diet and ketogenic diet. For each amino acid, two metrics are used to quantify their enrichment in a diet: one is the absolute daily intake of this amino acid in a diet, the other one is the fraction of this amino acid in the total daily intake of all amino acids. As we have discussed previously, there are 18 variables describing absolute levels of amino acids in foods and diets. These variables are the amounts of serine, tyrosine, glycine, phenylalanine, proline, valine, lysine, leucine, isoleucine, tryptophan, arginine, methionine, histidine, threonine, alanine, cystine, aspartate + asparagine, and glutamate + glutamine. The units are gram amino acid per gram weight of food (for amino acid levels in foods) and gram amino acid per day (for amino acid levels in diets).

#### 2.1 Calculation of ranges of amino acid levels in diets

As we have discussed in section 1.2, the absolute levels of amino acids in a diet $\boldsymbol{x}$ are $\boldsymbol{s} = \mathbf{A}\boldsymbol{x}$, and the absolute level of the $i$-th amino acid in this diet is $s_i = \boldsymbol{a}_i^T \boldsymbol{x}$. Hence, with the mathematical definition of a dietary pattern, which we have developed in the previous section, the lowest allowed intake of the $i$-th amino acid in this dietary pattern can be computed by solving the linear programming problem:

$$\min \boldsymbol{a}_i^T \boldsymbol{x}, \text{ s.t.}$$

$$\begin{cases} \boldsymbol{x} \geq \mathbf{0} \\ \boldsymbol{l}_n \leq \mathbf{C}\boldsymbol{x} \leq \boldsymbol{u}_n \\ \boldsymbol{l}_f \leq \mathbf{D}\boldsymbol{x} \leq \boldsymbol{u}_f \\ \mathbf{E}\boldsymbol{x} \leq \mathbf{0} \end{cases}$$

And the highest allowed intake of the i-th amino acid in this dietary pattern can be computed by solving another linear programming problem:

$$\max \boldsymbol{a}_i^T \boldsymbol{x}, \text{ s.t.}$$

$$\begin{cases} \boldsymbol{x} \geq \boldsymbol{0} \\ \boldsymbol{l}_n \leq \mathbf{C}\boldsymbol{x} \leq \boldsymbol{u}_n \\ \boldsymbol{l}_f \leq \mathbf{D}\boldsymbol{x} \leq \boldsymbol{u}_f \\ \mathbf{E}\boldsymbol{x} \leq \boldsymbol{0} \end{cases}$$

### 2.2 Sampling of amino acid composition in diets

Amino acid composition of a diet, or relative levels of amino acids in a diet, is defined as a vector $\boldsymbol{r}$ that contains ratios of absolute levels of amino acids to total amount of all amino acids in this diet:

$$r_i = \frac{s_i}{\sum_{j=1}^{18} s_j} = \frac{\boldsymbol{a}_i^T \boldsymbol{x}}{\sum_{j=1}^{18} \boldsymbol{a}_j^T \boldsymbol{x}} = \frac{\boldsymbol{a}_i^T \boldsymbol{x}}{[1 \quad \cdots \quad 1] \cdot \mathbf{A}\boldsymbol{x}}$$

Since this is no longer a linear function of $\boldsymbol{x}$, ranges of $r_i$ cannot be computed using linear

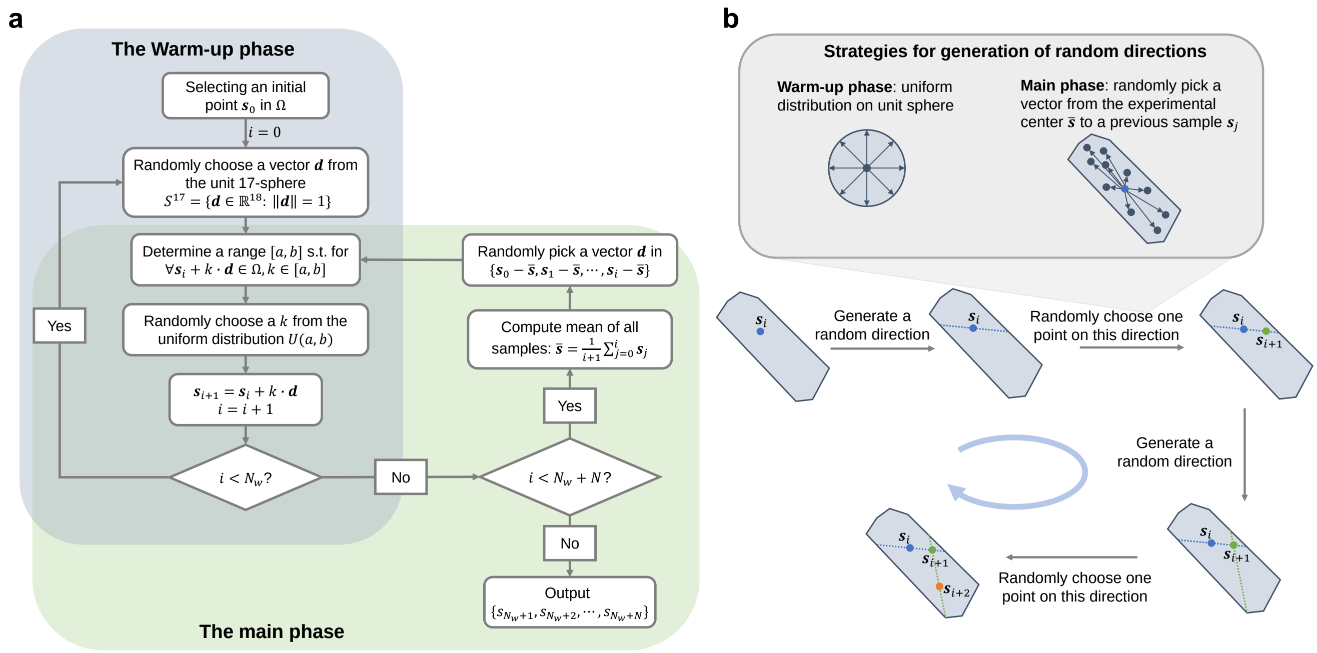

**Appendix Figure 1. The hit-and-run sampling algorithm.** (a) Workflow of the hit-and-run algorithm for uniform sampling of the feasible region for the amino acid abundance vector $\boldsymbol{s} = \mathbf{A}\boldsymbol{x}$. (b) Illustration of each iterative step and strategies for generation of random directions in the hit-and-run algorithm.

programming. Instead, we first uniformly sampled absolute levels of amino acids in all diets consistent with this dietary pattern (i.e. diets that satisfy all linear constraints set by definition of this dietary pattern):

$$\text{Uniformly sample } \boldsymbol{s} = \mathbf{A}\boldsymbol{x}, \text{ s.t.}$$

$$\begin{cases} \boldsymbol{x} \geq \mathbf{0} \\ \boldsymbol{l}_n \leq \mathbf{C}\boldsymbol{x} \leq \boldsymbol{u}_n \\ \boldsymbol{l}_f \leq \mathbf{D}\boldsymbol{x} \leq \boldsymbol{u}_f \\ \mathbf{E}\boldsymbol{x} \leq \mathbf{0} \end{cases}$$

Because all constraints on $\boldsymbol{x}$ are linear constraints, the feasible region for $\boldsymbol{x}$, i.e. the region in which a diet $\boldsymbol{x}$ falls in if and only if $\boldsymbol{x}$ is a diet consistent with this certain dietary pattern, is a polyhedron. Thus, because a polyhedron after a linear transformation is still a polyhedron, the feasible region for $\mathbf{A}\boldsymbol{x}$ is also a polyhedron. Let $\Omega$ denote this polyhedron. It was then uniformly sampled using the hit-and-run sampling algorithm with a direction choice strategy for accelerated convergence (ACHR, Appendix Figure 1)[1], generating $N$ samples $\boldsymbol{s}_1, \boldsymbol{s}_2, \cdots, \boldsymbol{s}_N$. These samples were then transformed to relative ratios of amino acids by the dividing total amount of amino acids in each sample:

$$\boldsymbol{r}_n = \frac{\boldsymbol{s}_n}{\sum_{i=1}^{18} s_{ni}}, n = 1,2,\cdots,N$$

In each iteration of the ACHR algorithm, a direction for perturbation, $\boldsymbol{d}$, is selected using different strategies based on the current phase of sampling (i.e. warm-up phase or main phase, Appendix Figure 1a), and the new sample $\boldsymbol{s}_{i+1}$ is generated by randomly perturb the current sample $\boldsymbol{s}_i$ on the direction $\boldsymbol{d}$ by the step size $k$ (Appendix Figure 1a):

$$\boldsymbol{s}_{i+1} = \boldsymbol{s}_i + k \cdot \boldsymbol{d}$$

Since the new sample $\boldsymbol{s}_{i+1}$ still needs to be in the feasible region, we have:

$$\begin{cases} \boldsymbol{s}_i + k \cdot \boldsymbol{d} = \mathbf{A}\boldsymbol{x} \\ \boldsymbol{x} \geq \mathbf{0} \\ \boldsymbol{l}_n \leq \mathbf{C}\boldsymbol{x} \leq \boldsymbol{u}_n \\ \boldsymbol{l}_f \leq \mathbf{D}\boldsymbol{x} \leq \boldsymbol{u}_f \\ \mathbf{E}\boldsymbol{x} \leq \mathbf{0} \end{cases}$$

Thus, the minimal and maximal allowed values for $k$ can be determined by solving the linear programming problems:

$$\min(\text{or max})\ k,\ \text{s.t.}$$

$$\begin{cases} [\mathbf{A} \quad -\boldsymbol{d}] \begin{bmatrix} \boldsymbol{x} \\ k \end{bmatrix} = \boldsymbol{s}_i \\ \boldsymbol{x} \geq \mathbf{0} \\ \boldsymbol{l}_n \leq \mathbf{C}\boldsymbol{x} \leq \boldsymbol{u}_n \\ \boldsymbol{l}_f \leq \mathbf{D}\boldsymbol{x} \leq \boldsymbol{u}_f \\ \mathbf{E}\boldsymbol{x} \leq \mathbf{0} \end{cases}$$

Let $[a, b]$ denote the range of $k$ determined by solving these two linear programming problems, we then randomly choose the value of $k$ from the uniform distribution on $[a, b]$ and generate the new sample $\boldsymbol{s}_{i+1}$ based on the randomly selected $k$:

$$k \sim U(a, b)$$

$$\boldsymbol{s}_{i+1} = \boldsymbol{s}_i + k \cdot \boldsymbol{d}$$

To determine the number $N$ of samples sufficient to thoroughly capture the distribution of amino acid compositions in a dietary pattern, we performed the ACHR sampling using different sample sizes for the USDA-recommended diet and examined the trends of mean and standard deviation against number of samples. We used the USDA-recommended diet here to determine $N$ because the definition of this dietary pattern includes the largest number of constraints among all dietary patterns studied here, thus likely exhibiting higher complexity in the sampling process compared to other dietary patterns. We found that for all amino acids, the mean and standard deviation become largely unchanged by increasing $N$ when $N$ is larger than 40,000. Thus, we used $N = 50{,}000$ to sample all dietary patterns. All linear programming problems were solved using the function 'mosekopt' in the MOSEK Optimization Toolbox for MATLAB[2].

### 3. Reconstruction of amino acid profiles in human diets

#### 3.1 Acquisition of datasets

Microsoft Access database files for USDA National Nutrient Database for Standard Reference (SR) and the USDA Food and Nutrient Database for Dietary Studies (FNDDS) were downloaded from the website for USDA Agricultural Research Service: https://www.ars.usda.gov/northeast-area/beltsville-md-bhnrc/beltsville-human-nutrition-research-center/methods-and-application-of-food-composition-laboratory/mafcl-site-pages/sr17-sr28/ (SR), and https://www.ars.usda.gov/northeast-area/beltsville-md-bhnrc/beltsville-human-nutrition-research-center/food-surveys-research-group/docs/fndds-download-databases/ (FNDDS). Matching between versions of the SR and FNDDS databases is shown in Appendix Table 1:

**Appendix Table 1.** Matching between versions of FNDDS and SR databases

| Year | FNDDS database version | SR database version |
|---|---|---|
| **2007-2008** | FNDDS 2007-2008 | Release 22 |
| **2009-2010** | FNDDS 2009-2010 | Release 24 |
| **2011-2012** | FNDDS 2011-2012 | Release 26 |
| **2013-2014** | FNDDS 2013-2014 | Release 28 |

Nutritional composition values of foods in the SR databases were retrieved from the table 'NUT_DATA' in the corresponding MS Access database files for the SR databases. Mappings from foods in the SR databases to foods in the FNDDS databases were retrieved from the table 'FNDDSSRLinks' in the MS Access database files for the FNDDS databases. Factors for moisture and fat adjustments (i.e. factors quantifying the amount of water or fat lost or gained during the cooking or food preparation processes for some foods in the database) were retrieved from the table 'MoistNFatAdjust' in the FNDDS database files. Retention factors for nutrients (i.e. factors quantifying the fraction of nutrients kept during cooking process) were obtained from the USDA Ag Data Commons website: https://data.nal.usda.gov/dataset/usda-table-nutrient-retention-factors-release-6-2007.

SAS (.xpt) files for NHANES 2007-2008, 2009-2010, 2011-2012, 2013-2014 datasets, including demographics data, dietary data, examination data, laboratory data and questionnaire data were retrieved from the web page for NHANES: https://wwwn.cdc.gov/nchs/nhanes/Default.aspx, and converted to R data frames using the function 'sasxport.get()' in the R package 'Hmisc'.

#### 3.2 Comparison of data imputation methods

Two methods for missing data imputation, random forest (RF) and predictive mean matching (PMM), were applied to the USDA SR Release 28 dataset to evaluate their performance. RF-based imputation was performed using the function 'missForest()' in the R package 'missForest'[3]. PMM-based imputation was performed using the function 'mice()' in the R package 'MICE'[4]. For each nutrient variable, we first calculated a missing ratio defined by the number of missing values for this variable divided by the total number of foods (i.e. 8788) in the dataset. All nutrient variables with missing ratio higher than 0.6 were discarded before the following analysis. We then adopted two alternative strategies for data transformation before the imputation. In one strategy (without transformation), the raw abundances of nutrients, including amino acids, were used as the input to the data imputation algorithms. In the other strategy, absolute abundances of amino acids were transformed to the ratio of absolute amino acid levels to one plus the protein level:

$$\widehat{Y_{\mathrm{AA}}} = \frac{Y_{\mathrm{AA}}}{1 + Y_{\mathrm{protein}}}$$

The transformed amino acid variables were then used together with raw values of other nutrient variables as the input to the data imputation algorithms. The major goal of performing this transformation is to decouple the variation in relative levels of amino acids and variation in absolute abundances of protein. After imputation has been done for the transformed variables $\widehat{Y_{\mathrm{AA}}}$, imputation for absolute levels of amino acids can be calculated from the imputed $\widehat{Y_{\mathrm{AA}}}$ values:

$$Y_{\mathrm{AA}}^{(\mathrm{i})} = \widehat{Y_{\mathrm{AA}}^{(\mathrm{i})}}\left(1 + Y_{\mathrm{protein}}^{(\mathrm{i})}\right)$$

In which the superscript (i) indicates imputed variables.

Combining the two imputation algorithms (i.e. RF and PMM) and two data transformation strategies, four data imputation scenarios were considered in the comparison. These include RF without data transformation, RF with data transformation, PMM without data transformation, and PMM with data transformation. Performance of these data imputation scenarios were evaluated using 5-fold cross validation, in which all values (including known values and missing values) in the input dataset were split into five groups. In the *n*-th (n=1,2,3,4,5) iteration, values in the *n*-th group were treated as the missing values which were then imputed using data points in other groups. After all five iterations were completed, the actual values and imputed values were compared for each nutrient using Spearman's rank correlation coefficient. Distributions of these Spearman's rank correlation coefficients were then compared across data imputation scenarios to determine the strategy with best performance (i.e. data imputation scenario that reached the highest Spearman correlation between the imputed and actual values). Spearman correlation coefficients between actual and imputed values for pairwise ratios between amino acid abundances (e.g. ratio of serine level to glycine level, etc) were also computed to evaluate the abilities of the data imputation methods to correctly capture relative abundances of amino acids in addition to their absolute levels. Since RF with data transformation showed the highest Spearman correlation coefficients between the actual and imputed values for both single nutrients (Supplementary Figure 4b) and pairwise ratios for amino acid abundances (Supplementary Figure 4c), this strategy was selected for imputation of all USDA SR datasets and FNDDS datasets.

#### 3.3 Imputation of USDA SR and FNDDS datasets

Missing data imputation was first done for nutritional composition data of foods in the USDA SR datasets. Nutrients with missing values in more than 60% of the foods were removed from the datasets before imputation. Amino acid abundances were transformed by dividing one plus the protein abundance as described in the previous section. Imputation was performed using the random forest regression method implemented in the function 'missForest()' in the R package 'missForest' with default parameters. Nutritional composition values of foods in the FNDDS datasets were computed using the imputed USDA SR datasets, the mapping information from foods in SR to foods in FNDDS, factors for moisture and fat adjustments, and retention factors for nutrients.

A typical food in the FNDDS dataset is prepared from one or more foods in the USDA SR dataset with the proportion for each food is provided together with its identifier in USDA SR. For instance, the FNDDS food with identifier '27520130' and description 'Bacon, chicken, and tomato club sandwich, with lettuce and spread' is mapped to six foods in the USDA SR database:

**Appendix Table 2**. Mapping between the USDA SR database and the food 'Bacon, chicken and tomato club sandwich' in the USDA FNDDS dataset

| USDA SR ID | Food name | Weight (g) |
|---|---|---|
| **18069** | Bread, white, commercially prepared (includes soft bread crumbs) | 78 |
| **5064** | Chicken, broilers or fryers, breast, meat only, cooked, roasted | 85 |
| **10124** | Pork, cured, bacon, cooked, broiled, pan-fried or roasted | 19 |
| **11252** | Lettuce, iceberg (includes crisphead types), raw | 25 |
| **11529** | Tomatoes, red, ripe, raw, year round average | 30 |
| **4025** | Salad dressing, mayonnaise, soybean oil, with salt | 13.75 |

Assuming that there are *m* foods in the USDA FNDDS database and *n* foods in the USDA SR database, we can define a $m \times n$ matrix $\mathbf{M}$ to describe the mapping from USDA SR to USDA FNDDS, in which the element $M_{ij}$ describes the amount of the *j*-th food in the USDA SR database needed in the preparation of the *i*-th food in the USDA FNDDS database. Let $\mathbf{Y}$ denote the nutritional composition of foods (i.e. amounts of nutrients in 1 gram of each food) in the USDA SR database, we can then compute the nutritional composition of the *i*-th food in the USDA FNDDS database:

$$\boldsymbol{z}_i = \frac{\sum_{j=1}^{n} M_{ij}\mathbf{R}_{ij}\boldsymbol{y}_j}{\sum_{j=1}^{n} M_{ij}}$$

in which $\mathbf{R}_{ij}$ is a diagonal matrix with the *k*-th diagonal element being the retention factor for the *k*-th nutrient in preparation of the *i*-th FNDDS food from the *j*-th SR food. A retention factor of 0.1 means that 10% of this nutrient is preserved during the food preparation process while the rest 90% is lost. This nutritional composition vector $\boldsymbol{z}_i$ was then further corrected for moisture and fat adjustments:

$$z_{ij}^{\text{(adjusted)}} = \begin{cases} \dfrac{100 \times z_{ij}}{100 + a^{(m)} + \sum_{k=1}^{n_f} a_k^{(f)}} & \text{if the } j\text{-th nutrient is neither water nor fat} \\ \dfrac{100 \times z_{ij} + a^{(m)}}{100 + a^{(m)} + \sum_{k=1}^{n_f} a_k^{(f)}} & \text{if the } j\text{-th nutrient is water} \\ \dfrac{100 \times z_{ij} + a_k^{(f)}}{100 + a^{(m)} + \sum_{k=1}^{n_f} a_k^{(f)}} & \text{if the } j\text{-th nutrient is the } k\text{-th fat adjusted} \end{cases}$$

in which $a^{(m)}$ is the moisture adjustment factor which quantifies grams of water gained or lost during cooking of 100 grams of raw food, $n_f$ is the total number of different types of fat, $\sum_{k=1}^{n_f} a_k^{(f)}$ is the total fat adjustment factor that quantifies total grams of fat gained or lost during cooking of 100 grams of raw food, $a_k^{(f)}$ is the fat adjustment factor that quantifies grams of the *k*-th type of fat (e.g. polyunsaturated fat, saturated fat 16:0, etc) gained or lost during cooking of 100 grams of raw food. The adjusted nutritional values for foods in FNDDS, including amino acid abundances in all FNDDS foods which were not included in the original FNDDS datasets, were used in the following reconstruction of amino acid intake profiles in the NHANES dietary records.

**3.4 Reconstruction of amino acid intake profiles in NHANES dietary records**

Nutrient composition values for dietary records in NHANES 2007-2008, 2009-2010, 2011-2012 and 2013-2014 datasets were computed based on the reconstructed nutritional values for foods in FNDDS and food consumption records in human subjects in the NHANES datasets (i.e. data files with the description ‘Dietary Interview – Individual Foods, First Day/Second Day’) retrieved from the web pages of NHANES. For each human subject, the food consumption record on one day (day 1 or day 2) includes a group of foods in FNDDS and grams of each food consumed on that day. An example of a food consumption record is shown in Appendix Table 3 below:

**Appendix Table 3**. An example of food consumption record in NHANES

| **seqn** (Identifier of human subject) | **dr1ifdcd** (FNDDS food ID) | **dr1igrms** (Grams of food) |
|---|---|---|
| 41475 | 64132010 | 160 |
| 41475 | 92101500 | 710.4 |
| 41475 | 91200040 | 4 |
| 41475 | 12120100 | 10.08 |
| 41475 | 52215200 | 63.62 |
| 41475 | 22600100 | 10 |
| 41475 | 71000100 | 20.13 |
| 41475 | 72201200 | 30 |
| 41475 | 75221000 | 4.48 |
| 41475 | 32104900 | 122 |
| 41475 | 94000100 | 474 |
| 41475 | 92302300 | 1554 |
| 41475 | 94000100 | 474 |
| 41475 | 27214110 | 134.3 |
| 41475 | 58145110 | 182.25 |
| 41475 | 75216123 | 42.25 |
| 41475 | 92302300 | 473.6 |
| 41475 | 53410100 | 122.06 |
| 41475 | 13110100 | 16.63 |
| 41475 | 94000100 | 59.25 |

Let **Z** denote the reconstructed nutritional values (which include the reconstructed amino acid abundances in these foods) for foods in FNDDS and $\boldsymbol{x}$ denote a food consumption vector in which the i-th value quantifies the total grams of the i-th food consumed on that day, we can compute the nutrient intake profile, which includes the uptake of amino acids, in this dietary record:

$$\boldsymbol{y} = \frac{\mathbf{Z} \cdot \boldsymbol{x}}{\|\boldsymbol{x}\|_1}$$

in which $\|\boldsymbol{x}\|_1$ is the L1-norm of the food consumption vector (i.e. total weight of food consumed on that day).

### 4. Analysis of association between dietary amino acids and human health

#### 4.1 Definition of human health variables

Binary disease variables indicating the presence of pathological conditions, including obesity, hypertension, diabetes, and cancer, were constructed based on the datasets 'Examination data', 'Laboratory data', and 'Questionnaire data' in the NHANES databases 2007-2008, 2009-2010, 2011-2012, and 2013-2014. Adults with BMI values higher than 30 were considered obese. Hypertension was defined as the condition of systolic blood pressure (the variables 'bpxsy1', 'bpxsy2' and 'bpxsy3', corresponding to three consecutive measurements) being higher than 120 mm Hg and diastolic blood pressure (the variables 'bpxdi1', 'bpxdi2' and 'bpxdi3', corresponding to three consecutive measurements) being higher than 80 mm Hg. Diabetes was defined as the condition of glycohemoglobin levels (the variable 'lbxgh') being higher than 6.5%, fasting plasma glucose concentration (the variable 'lbxglu') higher than 126 mg/dL, and blood glucose concentration in response to oral glucose tolerance test (the variable 'lbxglt') higher than 200 mg/dL. Information about the presence of cancer was obtained from answers to the question 'Have you ever been told by a doctor or other health professional that you had cancer or a malignancy of any kind?' in the questionnaire about medical conditions, in which the answers 'yes' or 'no' were linked to the presence or absence of cancer, while the answers 'refused' and 'don't know' were considered as missing data.

#### 4.2 Correlation analysis

Partial Spearman correlation coefficients between dietary amino acid composition or other nutritional variables and health variables defined in the previous section were computed using the MATLAB function ‘partialcorr’, controlling for demographic and lifestyle-related factors including income, education, age, gender, ethnicity, marital status, smoking, alcohol consumption, physical activity, and batch. Only adults (i.e. age > 20 years old) were included in the analysis. Individuals with dietary intake of any nutrient higher than three times of the 99$^{th}$ percentile of the intake of that nutrient among the population were considered outliers and not included in the following analysis.

**4.3 Machine learning**

Dietary variables were first adjusted to the demographic and lifestyle-related variables (i.e. potential confounding factors) by fitting linear regression models with the confounding factors as covariates and dietary variables as dependent variables. Let **X** denote the nutrient intake profiles in all individuals involved in the analysis (each column represents a nutrient and each row represents an individual) and **Z** for the confounding variables (each column represents a variable and each row represents an individual), we assume that the relationship between the intake of each specific nutrient and the demographic and lifestyle-related variables is linear:

$$\boldsymbol{X}_i = \boldsymbol{Z} \cdot \boldsymbol{c}_i + \boldsymbol{r}_i$$

In which $\boldsymbol{X}_i$ is the i-th column of the matrix **X** (i.e. intake profile of the i-th nutrient), $\boldsymbol{c}_i$ is the vector of linear coefficients, $\boldsymbol{r}_i$ is the residue term which represents for the part of intake of the i-th nutrient whose variance cannot be explained by variance in the confounding variables. Hence, the variables $\boldsymbol{r}_i$ were termed adjusted nutritional variables, meaning that they have been adjusted for the confounding variables. We next grouped the adjusted nutritional variables into seven categories: energy, vitamins, minerals, macronutrients, macronutrient compositions, amino acid compositions, other nutrients. The complete table of nutritional variables and the groups they belong to is included below.

**Appendix Table 4**. Seven categories of nutritional variables

| **Energy** | **Macronutrients** | **Macronutrient subtypes** | **Vitamins** | **Minerals** | **Amino acids** | **Others** |
|---|---|---|---|---|---|---|
| Energy(kc | Carbohydrate(g | PFA | Food | Zinc(mg) | Serine | Caffeine(mg) |

| | | | | | | |
|---|---|---|---|---|---|---|
| al) | m) | 18:2(Octadecadienoic)(gm) | folate(mcg) | | | |
| | Total fat(gm) | Total monounsaturated fatty acids(gm) | Vitamin K(mcg) | Iron(mg) | Tyrosine | Theobromine(mg) |
| | Protein(gm) | PFA 18:3(Octadecatrienoic)(gm) | Niacin(mg) | Copper(mg) | Glycine | |
| | | SFA 16:0(Hexadecanoic)(gm) | Riboflavin (Vitamin B2)(mg) | Phosphorus(mg) | Phenylalanine | |
| | | Dietary fiber(gm) | Folate,DFE(mcg) | Calcium(mg) | Proline | |
| | | Total sugars(gm) | Total choline(mg) | Sodium(mg) | Valine | |
| | | Total polyunsaturated fatty acids(gm) | Vitamin C(mg) | Selenium(mcg) | Lysine | |
| | | MFA 18:1 (Octadecenoic)(gm) | Vitamin A,RAE(mcg) | Magnesium(mg) | Aspartate+Asparagine | |
| | | SFA 18:0(Octadecanoic)(gm) | Vitamin B12(mcg) | Potassium(mg) | Leucine | |
| | | Total saturated fatty acids(gm) | Vitamin E as alpha-tocopherol(mg) | | Isoleucine | |

| | | | | | |
|---|---|---|---|---|---|
| | | | Thiamin(Vitamin B1)(mg) | | Tryptophan |
| | | | Vitamin B6(mg) | | Arginine |
| | | | Vitamin D(D2+D3)(mcg) | | Glutamate+Glutamine |
| | | | Folic acid(mcg) | | Methionine |
| | | | | | Histidine |
| | | | | | Threonine |
| | | | | | Alanine |
| | | | | | Cystine |

We let $\boldsymbol{R} = [\boldsymbol{r}_1, \boldsymbol{r}_2, \cdots, \boldsymbol{r}_k]$ be the matrix consisting of all adjusted nutritional variables in a category, and built a logistic regression model to predict the binary disease variables from the adjusted nutritional variables. The model was trained using the MATLAB function glmfit() for each category of nutritional variables, and the accuracy of its prediction was evaluated using the area under the receiver operating characteristic curve (AUC) with 5-fold cross validation, in which the entire dataset was split into five groups and the model was trained with four out of the five groups were used as training set while the remaining one group was used as the test set. Uncertainty in the accuracy of prediction was evaluated by repeating the cross validation procedure 10 times using different ways of splitting the dataset and calculating the mean and standard deviation of the AUC values.

### 5 Development of the diet designer

#### 5.1 Categories of association between dietary amino acid and obesity

For each amino acid, human adults included in the NHANES datasets were divided into five bins according to the quintiles of adjusted intake of that amino acid. Obesity incidence was then calculated for human subjects in each quintile by the fraction of people with obesity (defined as BMI > 30) in that quintile. Chi-squared test was performed to assess whether the obesity incidence differs across the five quintiles of amino acid intake. For an amino acid significantly associated with obesity incidence (i.e. Chi-squared p-value < 0.05), let $\boldsymbol{p} = [p_1, p_2, p_3, p_4, p_5]$ denote the obesity incidence in the 1[st] to 5[th] quintiles of the intake of this amino acid, we then determined the category that the amino acid-obesity association belongs to according to $\boldsymbol{p}$. If the minimal value of elements in $\boldsymbol{p}$ is $p_1$ and the maximal value is either $p_4$ or $p_5$, the amino acid was considered to be positively associated with obesity incidence. On the other hand, if the minimal value of elements in $\boldsymbol{p}$ is $p_5$ and the maximal value is either $p_1$ or $p_2$, the amino acid was considered negatively associated with obesity incidence. Otherwise, the amino acid was considered associated with obesity incidence with U-shaped relationship. The only exception was lysine, for which $\boldsymbol{p} = [0.3449 \quad 0.3786 \quad 0.3979 \quad 0.3917 \quad 0.3918]$, for which the U-shape relationship will be assigned according to the rules described previously, but since the values of $p_3, p_4$ and $p_5$ were close to each other and significantly higher than $p_1$ and $p_2$, this relationship is better described as a positive association. Hence, lysine was considered as an amino acid positively associated with obesity incidence.

#### 5.2 Analysis of Pareto optimality

Based on the amino acids positively or negatively associated with obesity incidence that were found in the previous section, we defined two objectives for optimization of dietary amino acid intake, that is, to maximize the total intake of amino acids negatively associated with obesity incidence (i.e. AAs-to-maximize), and to minimize the total intake of amino acids positively associated with obesity incidence (i.e. AAs-to-minimize). Let $\boldsymbol{a}_+$ and $\boldsymbol{a}_-$ denote the vectors consisting of total abundance of AAs-to-maximize and AAs-to-minimize in each food, then the inner products $\boldsymbol{a}_+^T\boldsymbol{x}$ and $\boldsymbol{a}_-^T\boldsymbol{x}$ indicate the total intake of AAs-to-maximize and AAs-to-minimize in a diet $\boldsymbol{x}$. Therefore, we have the mathematical form of optimizing the two amino acid intake goals for a specific dietary pattern with the general form described in section 1.3.4:

$$\max \boldsymbol{a}_+^T\boldsymbol{x}, \min \boldsymbol{a}_-^T\boldsymbol{x}, \text{s.t.}$$

$$\begin{cases} \boldsymbol{x} \geq \mathbf{0} \\ \boldsymbol{l}_n \leq \mathbf{C}\boldsymbol{x} \leq \boldsymbol{u}_n \\ \boldsymbol{l}_f \leq \mathbf{D}\boldsymbol{x} \leq \boldsymbol{u}_f \\ \mathbf{E}\boldsymbol{x} \leq \mathbf{0} \end{cases}$$

A feasible solution $\boldsymbol{x}_0$ of this problem is defined as a Pareto solution if for any other feasible solution $\boldsymbol{x}_1$, $\boldsymbol{a}_-^T\boldsymbol{x}_1 > \boldsymbol{a}_-^T\boldsymbol{x}_0$ if $\boldsymbol{a}_+^T\boldsymbol{x}_1 > \boldsymbol{a}_+^T\boldsymbol{x}_0$, and $\boldsymbol{a}_+^T\boldsymbol{x}_1 < \boldsymbol{a}_+^T\boldsymbol{x}_0$ if $\boldsymbol{a}_-^T\boldsymbol{x}_1 < \boldsymbol{a}_-^T\boldsymbol{x}_0$. In other words, for any other feasible solution $\boldsymbol{x}_1$, it is impossible that $\boldsymbol{x}_1$ has better performance than $\boldsymbol{x}_0$ in both of the two objectives of maximizing total AAs-to-maximize and minimizing total AAs-to-minimize. If the diet $\boldsymbol{x}_1$ has higher total intake of AAs-to-maximize than the diet $\boldsymbol{x}_0$, then it must have higher total intake of AAs-to-minimize than $\boldsymbol{x}_0$. On the other hand, if the diet $\boldsymbol{x}_1$ has lower total intake of AAs-to-minimize than the diet $\boldsymbol{x}_0$, then it must have lower total intake of AAs-to-maximize than $\boldsymbol{x}_0$. The Pareto surface of the problem is then defined as the set consisting of all Pareto solutions within the feasible region.

To construct the Pareto surface, we applied the $\varepsilon$-Constraint algorithm. Briefly, for a human dietary pattern with defined mathematical form, we first determine the range of total intake of AAs-to-maximize in that dietary pattern using linear programming, $[q, r]$:

$$u = \min \boldsymbol{a}_+^T\boldsymbol{x}, \text{s.t.}$$

$$\begin{cases} \boldsymbol{x} \geq \mathbf{0} \\ \boldsymbol{l}_n \leq \mathbf{C}\boldsymbol{x} \leq \boldsymbol{u}_n \\ \boldsymbol{l}_f \leq \mathbf{D}\boldsymbol{x} \leq \boldsymbol{u}_f \\ \mathbf{E}\boldsymbol{x} \leq \mathbf{0} \end{cases}$$

$$v = \max \boldsymbol{a}_+^T\boldsymbol{x}, \text{s.t.}$$

$$\begin{cases} \boldsymbol{x} \geq \mathbf{0} \\ \boldsymbol{l}_n \leq \mathbf{C}\boldsymbol{x} \leq \boldsymbol{u}_n \\ \boldsymbol{l}_f \leq \mathbf{D}\boldsymbol{x} \leq \boldsymbol{u}_f \\ \mathbf{E}\boldsymbol{x} \leq \mathbf{0} \end{cases}$$

Therefore, for any value $s \in [q, r]$, a Pareto solution can be obtained by solving the linear programming problem below:

$$\min \boldsymbol{a}_-^T\boldsymbol{x}\text{, s.t.}$$
$$\begin{cases} \boldsymbol{x} \geq \boldsymbol{0} \\ \boldsymbol{l}_n \leq \mathbf{C}\boldsymbol{x} \leq \boldsymbol{u}_n \\ \boldsymbol{l}_f \leq \mathbf{D}\boldsymbol{x} \leq \boldsymbol{u}_f \\ \mathbf{E}\boldsymbol{x} \leq \boldsymbol{0} \\ \boldsymbol{a}_+^T\boldsymbol{x} \geq w \end{cases}$$

The Pareto surface of each dietary pattern was constructed by uniformly selecting 100 values of $s \in [q, r]$ and computing the corresponding Pareto solution using the method described above.

#### 5.3 Estimation of deviation from Pareto surface

For each dietary record in the NHANES dataset, we first computed the total daily intake of AAs-to-maximize and AAs-to-minimize in that dietary record: $(u, v)$, and then computed the Euclidean distance between $(u, v)$ and the Pareto surface constructed using the method described in the previous section. Let $\boldsymbol{\Theta} = \{(s_i, t_i)\}_{i=1,2,\cdots,100}$ denote the set consisting of the intake of AAs-to-maximize and AAs-to-minimize in the 100 Pareto solutions computed previously, in which $s_i$ and $t_i$ means the intake of AAs-to-maximize and AAs-to-minimize in the i-th Pareto solution, respectively. We then calculated the deviation from Pareto surface:

$$d(u, v) = \min_{i=1,2,\cdots,100} \|(u, v) - (s_i, t_i)\|_2$$

Since the deviation from the Pareto surface computed this way is largely dependent of the total intake of protein in the dietary record, we adjusted it to the protein intake by fitting a 6-th order polynomial function of protein intake. Let $z$ denote the intake of protein in a dietary record (which has the intake of AAs-to-maximize and AAs-to-minimize being $(u, v)$, we fit the data to the model below:

$$d(u, v) = f(z) = c_0 + c_1 z + c_2 z^2 + c_3 z^3 + c_4 z^4 + c_5 z^5 + c_6 z^6$$

The residue $\hat{d} = d(u, v) - f(z)$ was then defined as the adjusted deviation from Pareto surface, which was used in all downstream analyses related to deviation from Pareto surface (Appendix Figure 2).

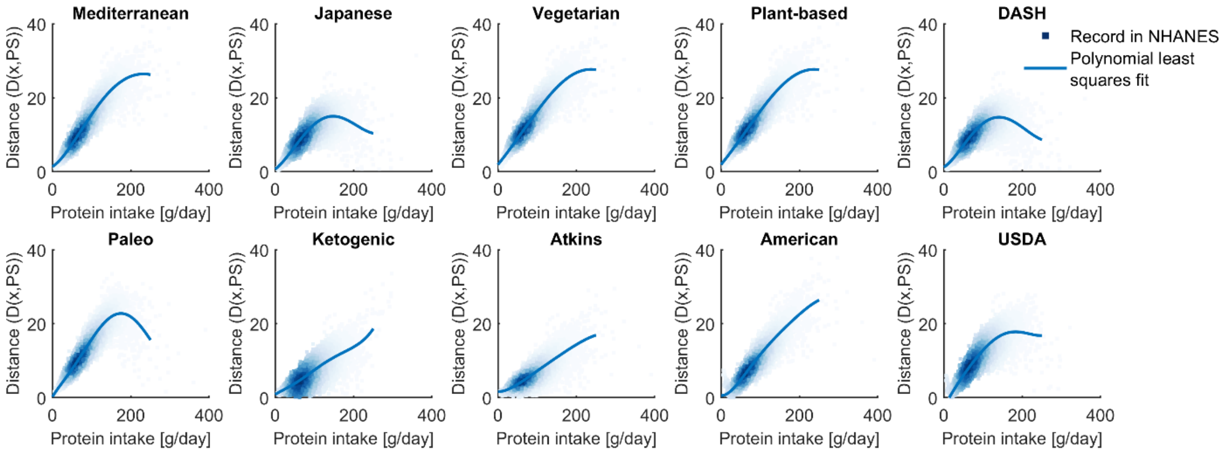

**Appendix Figure 2. Adjustment of the deviation from Pareto surface to total protein intake.**

1 Kaufman, D. E. & Smith, R. L. Direction choice for accelerated convergence in hit-and-run sampling. *Oper Res* **46**, 84-95, doi:DOI 10.1287/opre.46.1.84 (1998).

2 Andersen, E. D. & Andersen, K. D. in *High Performance Optimization* (eds Hans Frenk, Kees Roos, Tamás Terlaky, & Shuzhong Zhang) 197-232 (Springer US, 2000).

3 Stekhoven, D. J. & Buhlmann, P. MissForest-non-parametric missing value imputation for mixed-type data. *Bioinformatics* **28**, 112-118, doi:10.1093/bioinformatics/btr597 (2012).

4 van Buuren, S. & Groothuis-Oudshoorn, K. mice: Multivariate Imputation by Chained Equations in R. *J Stat Softw* **45**, 1-67 (2011).

### Supplementary Figures

#### Mediterranean diet

| Food | Servings |
|---|---|
| Cereals | 3~6/day |
| Fish | >=2/week |
| Fruits | 3~6/day |
| Legumes | >=2/week |
| White meat | 2/week |
| Processed meat | <=1/week |
| Sweets | <=2/week |
| Vegetables | >=6/day |
| Potatoes | <=3/week |
| Red meat | <=2/week |
| Olives/nuts/seeds | 1~2/day |
| Olive oil | >=1/day |
| Eggs | 2~4/week |
| Dairy | 2/day |
| Others | 0/day |

#### Paleo diet

| Food | Servings |
|---|---|
| Game meat/seafood | unlimited |
| Nuts/seeds | unlimited |
| Fruits/vegetables | unlimited |
| Honey | unlimited |
| Others | 0/day |

#### Vegetarian diet

| Food | Servings |
|---|---|
| Meat | 0/day |
| Others | unlimited |

#### Japanese diet

| Food | Servings |
|---|---|
| Cereals | 5~7/day |
| Fruits | 2/day |
| Vegetables | 5~6/day |
| Dairy | 2/day |
| Proteins | 3~5/day |
| Others | 0/day |

#### American diet

| Food | Consumption (+/-20%) |
|---|---|
| Meat | 381g/day |
| Sweets | 166g/day |
| Fats/oils | 114g/day |
| Grains | 291g/day |
| Starchy roots | 164g/day |
| Vegetables | 310g/day |
| Fruits | 266g/day |
| Eggs | 38g/day |
| Milk | 704g/day |
| Legumes | 9g/day |
| Alcoholic beverages | 258g/day |
| Others | 28g/day |
| **Nutrient** | **Intake (+/-20%)** |
| Calories | 3641 kcal/day |

#### Ketogenic diet

| Nutrient | % kcal |
|---|---|
| Fat | >=70% |
| Carbohydrate | <=5% |

#### USDA-recommended diet

| Nutrient | Intake |
|---|---|
| Calories | 1800~2200kcal/day |
| Protein | >=46g/day, 10~35%kcal |
| Carbohydrate | >=130g/day, 45~65%kcal |
| Dietary fiber | >=28g/day |
| Sugar | <=10%kcal |
| Fat | 20~35%kcal |
| Saturated fat | <=10%kcal |
| Linoleic acid | >=12g/day |
| Linolenic acid | >=1.1g/day |
| Calcium | >=1g/day |
| Iron | >=18mg/day |
| Magnesium | >=310mg/day |
| Phosphorus | >=700mg/day |
| Potassium | >=4.7g/day |
| Sodium | <=2.3g/day |
| Zinc | >=8mg/day |
| Copper | >=0.9mg/day |
| Manganese | >=1.8mg/day |
| Selenium | >=55μg/day |
| Vitamin A | >=700mg/day |
| Vitamin E | >=15mg/day |
| Vitamin D | >=600IU/day |
| Vitamin C | >=75mg/day |

#### Atkins diet

| Nutrient | Intake |
|---|---|
| Carbohydrate | <=20g/day |

#### USDA-recommended diet (continued)

| Nutrient | Intake |
|---|---|
| Thiamin | >=1.1mg/day |
| Riboflavin | >=1.1mg/day |
| Niacin | >=14mg/day |
| Vitamin B6 | >=1.3mg/day |
| Vitamin B12 | >=2.4μg/day |
| Choline | >=425mg/day |
| Vitamin K | >=90μg/day |
| Folate | >=400μg/day |

#### DASH diet

| Food | Servings |
|---|---|
| Grains | 6~8/day |
| Vegetables | 4~5/day |
| Fruits | 4~5/day |
| Low-fat dairy | 2~3/day |
| Lean meat | <=6/day |
| Nuts/seeds/legumes | 4~5/week |
| Fats/oils | 2~3/day |
| Sweets | <=5/week |
| Others | 0/day |
| **Nutrient** | **Intake** |
| Sodium | <=2.3g/day |

#### Plant-based diet

| Food | Servings |
|---|---|
| Meat | 0/day |
| Dairy | 0/day |
| Eggs | 0/day |
| Others | unlimited |

**Supplementary Figure 1 (Related to Figure 2). Definition of human dietary patterns.**

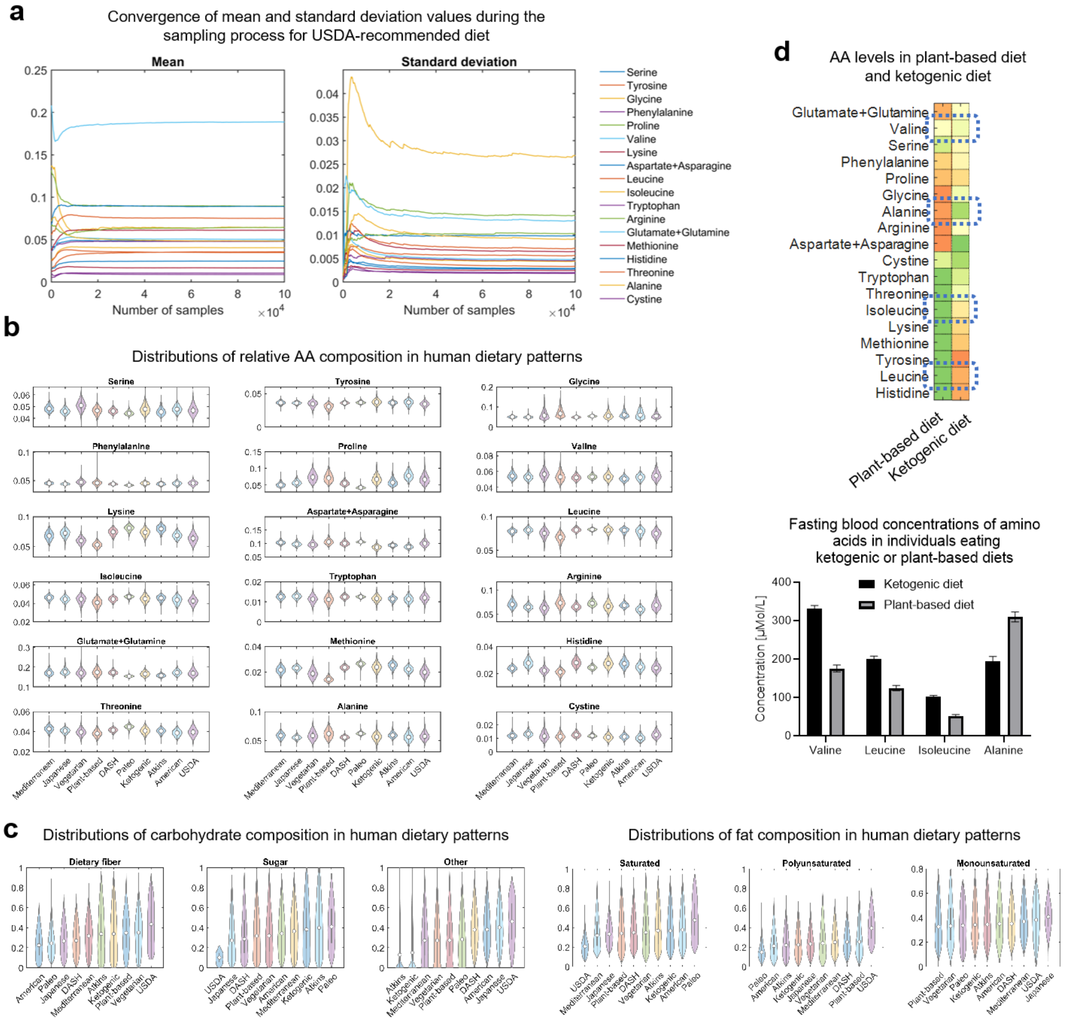

**Supplementary Figure 2 (Related to Figure 2). Amino acids, carbohydrates, and fats in human dietary patterns.**

(a) Convergence of mean and standard deviation values during the sampling of diets under the dietary pattern "USDA recommended diet".

(b) Distributions of relative amino acid composition in human dietary patterns.

(c) Distributions of relative carbohydrate composition and relative fat composition in human dietary patterns.

(d) Comparison between amino acid levels in plant-based and ketogenic diets estimated by our computational method and fasting blood amino acid concentrations in human individuals eating plant-based or ketogenic diet.

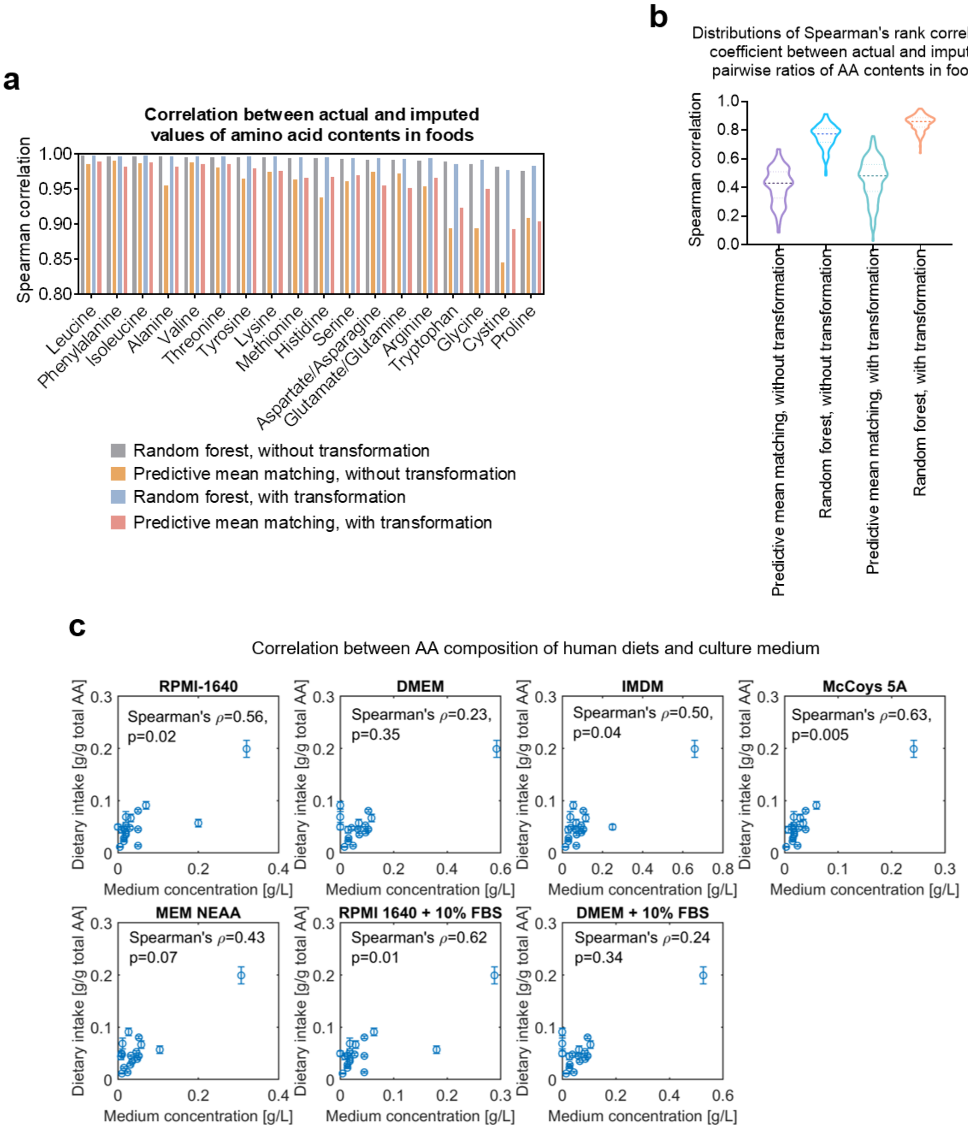

**Supplementary Figure 3 (Related to Figure 3). Imputation and validation of amino acid intake profiles in human dietary records**.

(a) Spearman's rank correlation coefficients between actual and imputed values of amino acid abundances in foods using different algorithms for data imputation.

(b) Distributions of Spearman's rank correlation coefficients between actual and imputed pairwise ratios of amino acid abundances in foods using different algorithms for data imputation.

(c) Comparison between concentrations of amino acids in culture mediums and imputed human dietary amino acid intakes.

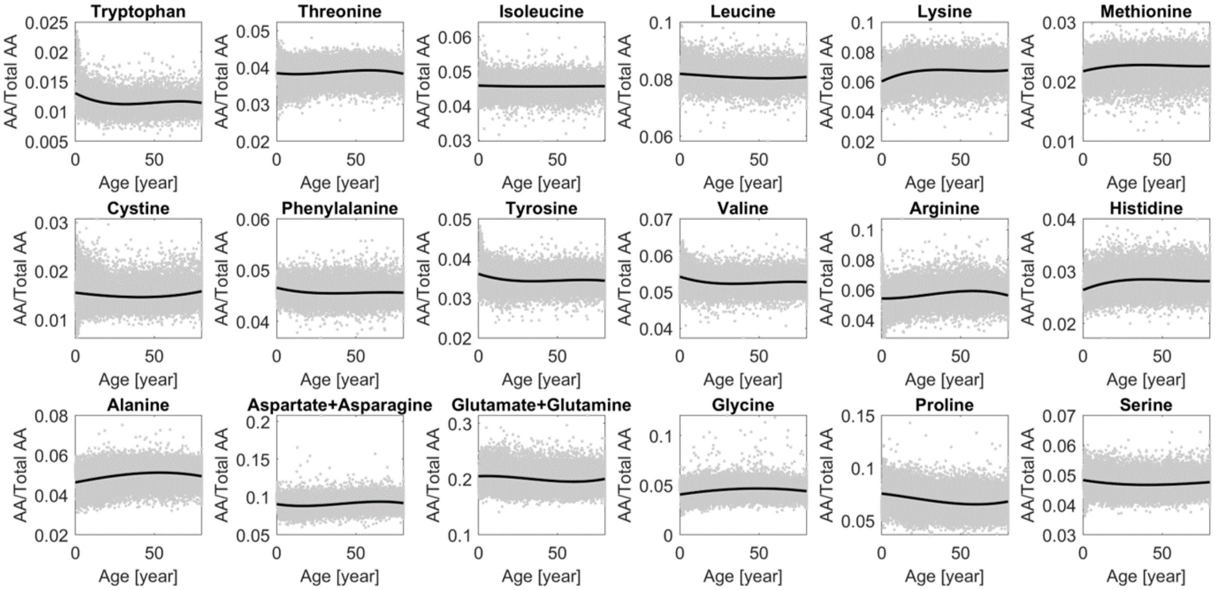

**Supplementary Figure 4 (Related to Figure 3). Relationship between dietary amino acid intake and age in U.S. citizens.**

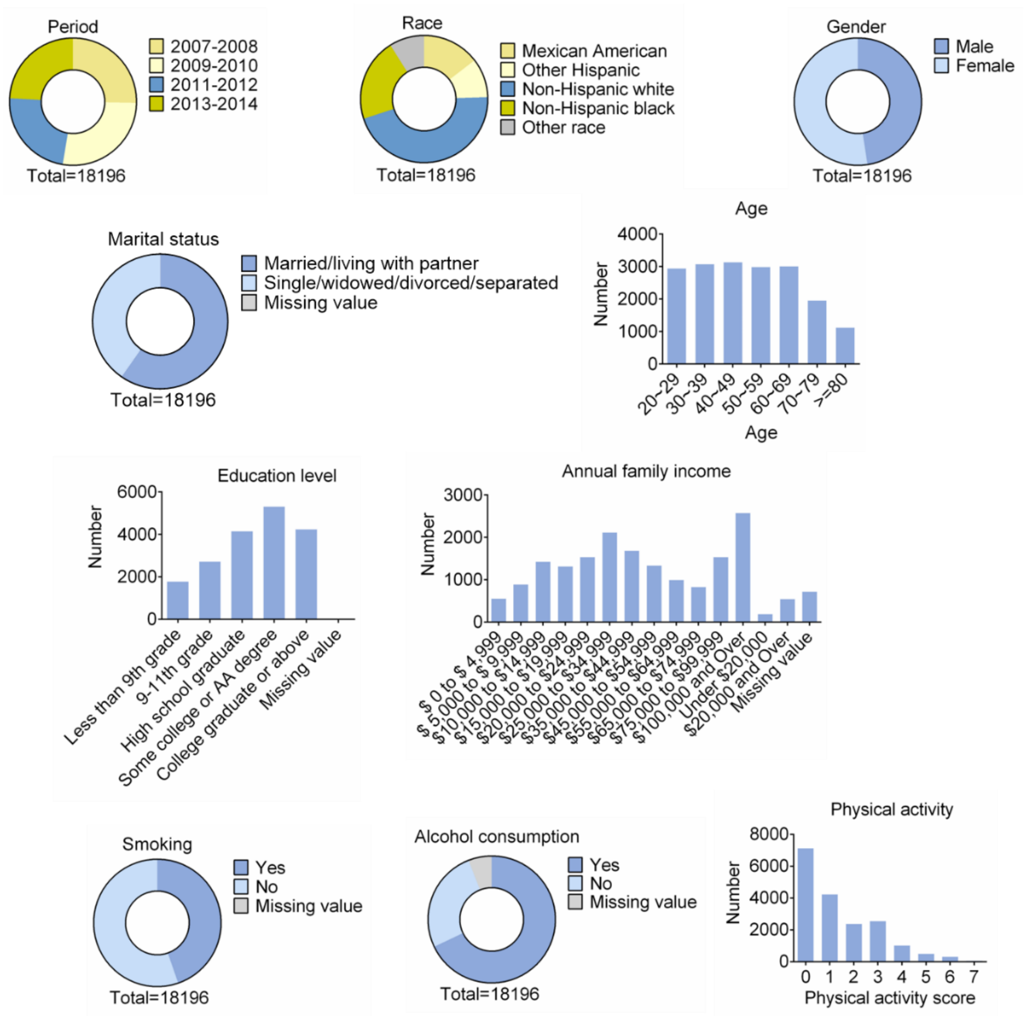

**Supplementary Figure 5 (Related to Figure 4). Demographic characteristics of the population.**

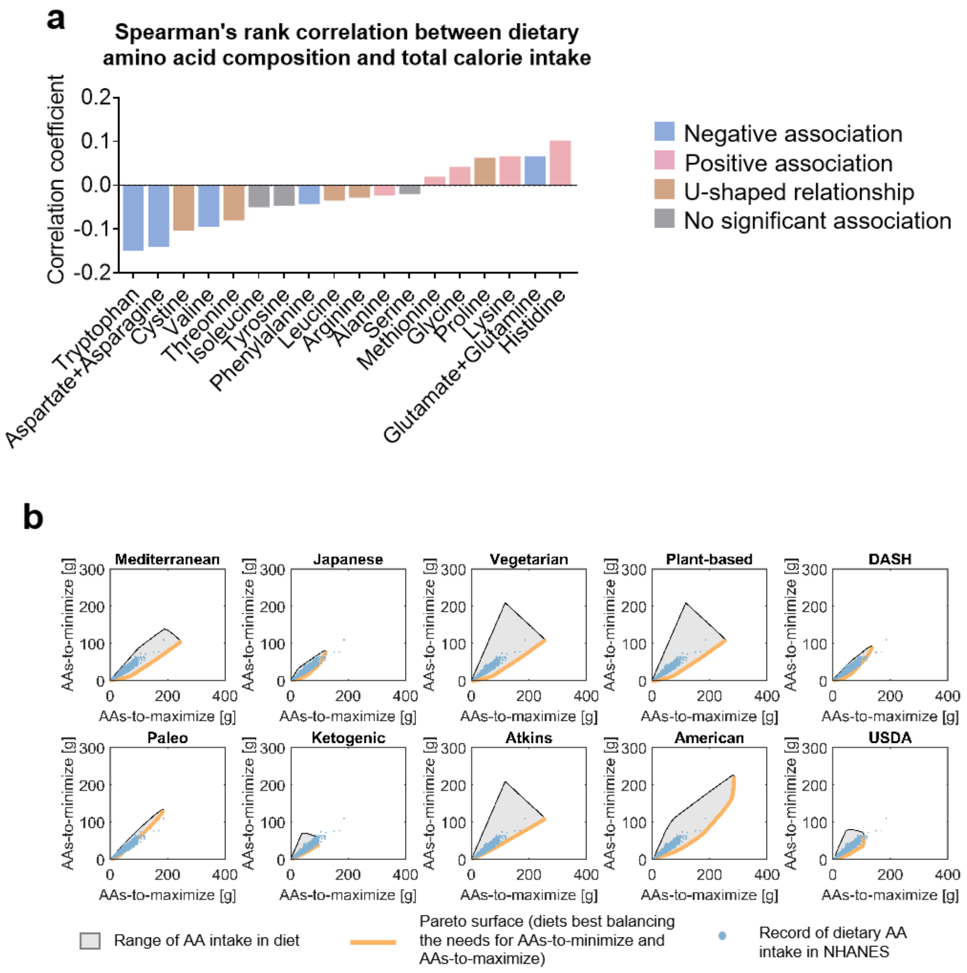

**Supplementary Figure 6 (Related to Figure 5). Association between dietary amino acid intake and obesity**.

(a) Spearman's rank correlation coefficients between dietary intake of amino acids and total calorie intake.

(b) Ranges of intake of total amino-acids-to-maximize and amino-acids-to-minimize in the 10 human dietary patterns (grey shaded regions), the Pareto surface (orange bold curve) corresponding to the two guidelines, i.e. maximizing total amino-acids-to-maximize, and minimizing total amino-acids-to-minimize, and actual intake of total amino-acids-to-maximize and total-amino-acids-to-minimize in human dietary records in the NHANES datasets.
